# Structural and Catalytic Architecture of the Cysteine Synthase Cys1a

**DOI:** 10.64898/2026.06.04.730285

**Authors:** Isis Pinner, Isabel N. Schultz, Jordan L. Pederick, Blagojce Jovcevski, Tara L. Pukala, Stephen G. Bell, John B. Bruning

## Abstract

Antifungal drug resistance in *Aspergillus fumigatus* is a major global health concern. Due to the increasing frequency of drug-resistant strains, there is a growing need for novel antifungal drugs. Inhibition of the cysteine biosynthesis pathway in *A. fumigatus* represents a promising strategy for the development of antifungal adjuvants. Within this pathway, the previously uncharacterised enzyme AfuCys1a fulfils several desirable characteristics as an adjuvant target. To enable structure-based inhibitor design, AfuCys1a was structurally and functionally characterised using X-ray crystallography, biochemical assays, and comparative sequence and structural analysis. In the search for effective and viable adjuvants, this research characterises the catalytic mechanism and substrate recognition of AfuCys1a and identifies 2-picolinic acid and related compounds as promising starting points for inhibitor development, working towards the larger goal of combating rising deaths from infection by *A. fumigatus*.

## Introduction

Antifungal resistance is a growing global health concern, threatening the effective treatment of an ever-increasing number of infections caused by fungal pathogens. Among these fungal threats, *Aspergillus fumigatus* (*A. fumigatus*), a ubiquitous, filamentous fungus found worldwide in soil and decaying organic matter, contributes substantially to this global crisis, being responsible for 70% of fungal-associated deaths (Bongomin et al., 2018). Its remarkable resilience to harsh environmental conditions has enabled this fungus to become a prevalent opportunistic pathogen. During its lifecycle, *A. fumigatus* produces conidia; these are asexually produced spores which can be rapidly aerosolised and inhaled by human individuals. Once inside the human lung, conidia exhibit robust properties that promote infection. While healthy individuals eliminate these spores by innate immune mechanisms, immunocompromised individuals lack the same defences and struggle to fight infection (Latgé & Chamilos, 2019; Liu et al., 2024; Rankin, 1953). The infection can then progress to a form of the disease known as invasive aspergillosis (IA), where hyphae penetrate the lung epithelium and disseminate into the bloodstream, spreading to various organs and tissues throughout the body (Baltussen et al., 2019; Chai et al., 2010; Rivera et al., 2006). Unfortunately, in the context of immune-compromised individuals, recent estimates suggest an IA-associated crude mortality rate of over 85% (Denning, 2024). Consequently, *A. fumigatus* has been declared a priority fungal pathogen by the World Health Organisation (WHO) (Fisher & Denning, 2023).

The primary treatment against IA are triazoles, such as voriconazole and isavuconazole, which have helped lower mortality rates (Morrissey et al., 2024). These drugs inhibit the enzyme Cyp51A and consequently disrupt maintenance of fungal cell walls (Burks et al., 2021; Camps et al., 2012). However, azole-resistant strains are now being reported globally, a consequence of poor antifungal management within agricultural and environmental settings (Burks et al., 2021; Camps et al., 2012; Chen et al., 2013; Rhodes et al., 2022; Sen et al., 2023). Genomic analysis identified both *cyp51A* and non-*cyp51A* genetic variants linked to drug resistance, suggesting mutations within several *A. fumigatus* genes contribute to reduced efficacy of triazoles targeting Cyp51A. This emphasises the importance of exploring novel antifungal drug targets as well as methods that exert less selective pressure toward drug resistance (Barber et al., 2021; Sen et al., 2024). Unfortunately, developing drugs against novel fungal targets is challenging. Like humans, fungi are eukaryotes and therefore share conservation across several biological pathways, increasing the risk of off-target toxicity in humans for many novel antifungal targets (Campoy & Adrio, 2017).

Amidst these challenges, targeting the cysteine biosynthesis pathway of *A. fumigatus* presents as an attractive strategy for novel antifungal development. Cysteine is a proteinogenic amino acid with a reactive sulfhydryl group that contributes to protein stability, enzyme catalysis, redox regulation, post-translational modifications, and sulphur homeostasis (Amich, 2024). There are two main pathways for the synthesis of L-cysteine in living organisms, the sulphur assimilation pathway and the reverse transsulfuration pathway. In general, mammals will exclusively use the reverse transsulfuration pathway, while plants and bacteria primarily use the sulphur assimilation pathway, and fungi are known to use either or both (Tao et al., 2024). The classical and most prevalent sulphur assimilation pathway involves conversion of L-serine into O-acetyl-L-serine (L-OAS) by an L-serine O-acetyltransferase (SAT). L-OAS is then converted into L-cysteine by an L-OAS sulfhydrylase (OASS). Recently, a novel sulphur assimilation pathway has been characterised in the yeast *Schizosaccharomyces pombe* and the coral *Acropora loripes*, where L-serine is converted into O-succinyl-L-serine (L-OSS) by an L-serine O-succinyltransferase (Cys2) and then into L-cysteine by L-OSS sulfhydrylase (Cys1a) (Bastard et al., 2017; Salazar et al., 2022).

For *S. pombe* and *A. loripes*, L-cysteine is produced exclusively by Cys1a through sulphur assimilation, while *A. fumigatus* can produce L-cysteine through either reverse transsulfuration or sulphur assimilation (Amich et al., 2016; Bastard et al., 2017; Salazar et al., 2022). For *A. fumigatus*, *mecA* encodes a cystathionine-β-synthase (AfuCBS) that functions within the reverse transsulfuration pathway, while *CysB* encodes Cys1a (AfuCys1a), representing two key enzymes from each cysteine biosynthesis pathway (Amich et al., 2016; Bastard et al., 2017; Salazar et al., 2022). While single AfuCBS or AfuCys1a mutants grow normally and are fully virulent, the double mutant results in cysteine auxotrophy for *A. fumigatus*, suggesting relatively low redundancy within this pathway. The double mutant also displayed reduced growth under cysteine-limiting conditions, abolished conidial germination, impaired recovery from nutrient starvation, and reduced virulence in murine models (Amich et al., 2016). This demonstrated that *A. fumigatus* cannot fully compensate for cysteine auxotrophy, and cysteine availability within murine lung tissue is not sufficient for full virulence; targeted disruption of cysteine biosynthesis could exploit a metabolic vulnerability unique to the site of infection (Amich, 2024; Amich et al., 2016). Lastly, the Cys1a pathway is widely present throughout the fungi kingdom, but is absent in humans and other vertebrates (Salazar et al., 2022). This would offer potential for the development of inhibitors that are highly specific to this pathway in *A. fumigatus*, lowering the risk of off-target effects towards humans (Fang et al., 2013, 2015; Hurtado-Guerrero et al., 2008; Mabanglo et al., 2014; Y. Zhang et al., 2020). Collectively, these findings underscore the important role of cysteine biosynthesis for the virulence and survival of *A. fumigatus* and identifies the AfuCys1a metabolic pathway as a promising target for development of novel antifungal drugs.

## Materials and Methods

**Materials and resources tables.**

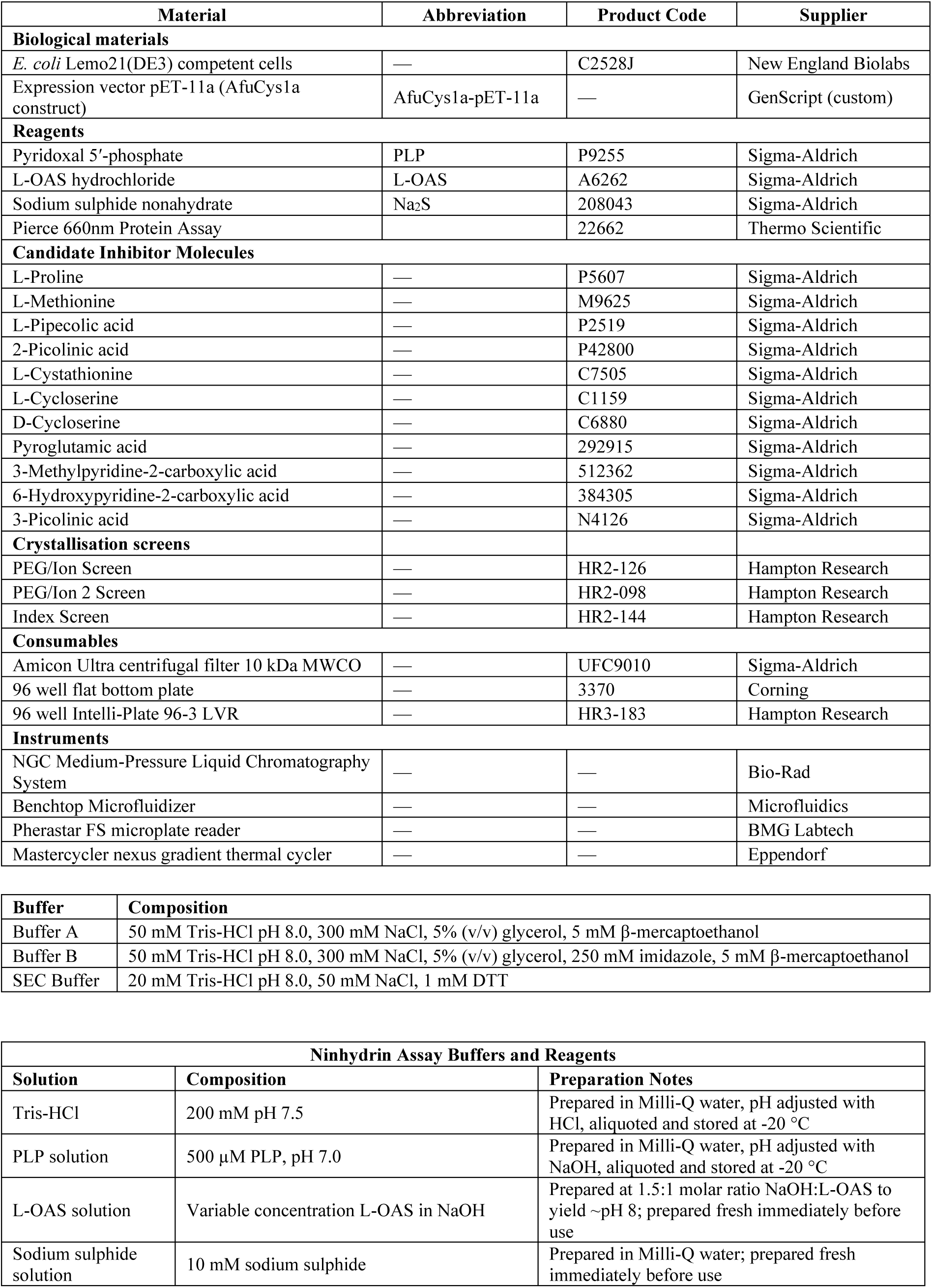
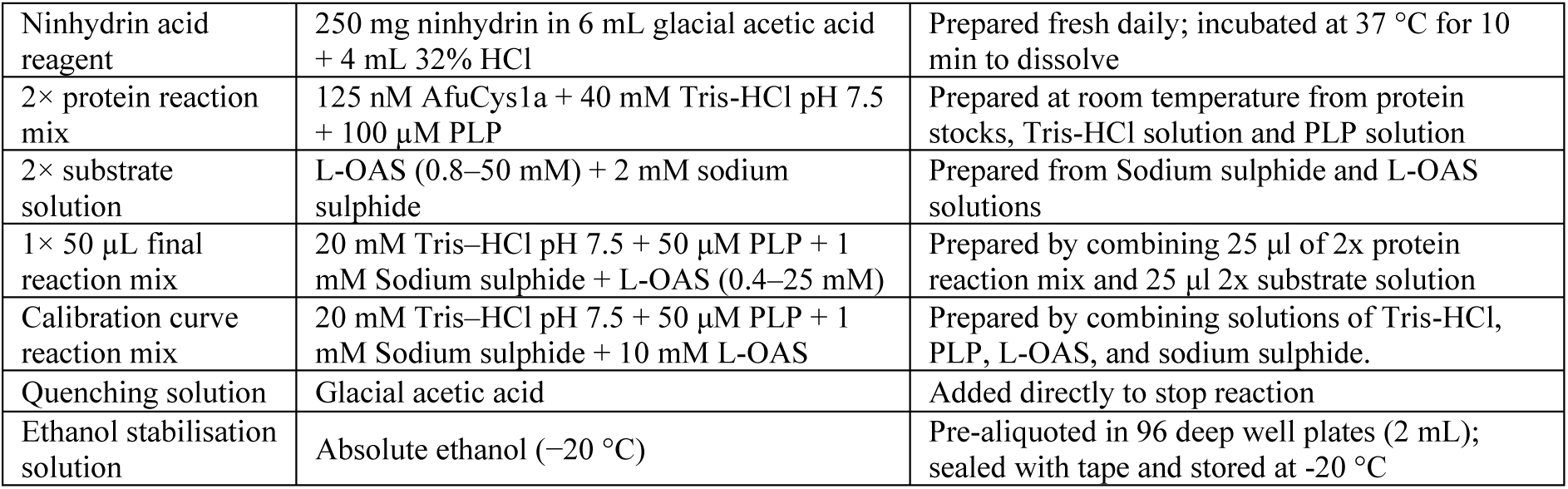

### Protein expression and purification

The pET-11a-AfuCys1a expression plasmid was synthesised by Genscript and contained the full-length AfuCys1a sequence (codon optimised for *Escherichia coli*), an N-terminal x6His tag, and a TEV cleavage site (ENLYFQS). The plasmid was transformed into *E. coli* Lemo21(DE3). Transformants were grown in LB media supplemented with 100 µg/mL ampicillin until OD_600_ = 0.5 (37°C, 180 rpm) and Pyridoxal 5′-phosphate (PLP) was added to a final concentration of 100 µM. Protein expression was induced by the addition of isopropyl-β-d-thiogalactopyranoside (IPTG) to a final concentration of 1 mM and continued for 16 hours at (16°C, 180 rpm).

Cells were pelleted by centrifugation at 5000 × *g* for 10 min at 4°C. Cell pellets were resuspended in Buffer A, lysed by mechanical disruption, and the lysate was clarified by centrifugation at 48,000 × *g* for 60 min at 4°C. The supernatant was applied to a Ni-NTA HisTrap HP column (equilibrated with Buffer A) and AfuCys1a was eluted with 60-100 % Buffer B. For size exclusion chromatography (SEC), elution fractions containing AfuCys1a were combined and concentrated using centrifugal filters (4°C, 4000 × *g*) to a final volume of 5 mL and applied to a HiPrep 26/60 Sephacryl S-200 HR gel filtration column (equilibrated with SEC buffer). This same buffer was used for elution at a flow rate of 0.5 mL per minute, which separated two distinct species of AfuCys1a; The first species was above 40 kDa and the second species was closer to 40 kDa. Fractions containing the second species (40 kDa) were combined and concentrated using centrifugal filters to 14.9 mg/mL, as determined by the pierce protein assay. A portion of the protein was used immediately for crystal screening trays while the remainder was stored at −80°C until further use. For enzymatic assays, a portion of AfuCys1a was thawed on ice and diluted in a solution containing 40 mM Tris-HCl pH 7.5 and 100 µM PLP to a final concentration of 10 µM. The diluted protein was then flash-frozen in liquid nitrogen and stored at −80°C.

### Crystallisation of AfuCys1a

Crystal screening trays were prepared with freshly purified AfuCys1a by the sitting drop vapour diffusion method, using PEG/Ion 1 & 2, and Index sparse matrix screens. Using 96-well Intelli-Plates, reservoirs contained 80 µL of each crystallisation reagent, with drops formed by adding 1 µL of AfuCys1a (14.9 mg/mL) with 1 µL of crystallisation reagent to the protein crystallisation well. Screening trays were stored at 16°C. The structures discussed here were produced from crystals that formed in well A2 and G2 of PEG/Ion 1 & 2, and G6 of Index. These structures are the1.47 Å LigA-bound and holo states, the 1.46 Å LigB-bound state, and the 1.60 Å acetate-bound state, respectively. All were a visible yellow colour, indicating presence of the PLP cofactor.

### X-ray Diffraction Data Collection, Processing, Phasing and Refinement

Crystals were immersed in a cryoprotectant consisting of 30% glycerol and 70% of the corresponding crystallisation reagent, and flash frozen in liquid nitrogen. Datasets were collected at the MX1 and MX2 beamlines of the Australian Synchrotron (Aragão et al., 2018; Cowieson et al., 2015). Indexing and integration were completed using XDS, with Aimless (CCP4) used for scaling and merging datasets (Kabsch, 2010; Winn et al., 2011). The AfuCys1a 1.47 Å LigA/holo structure was solved by molecular replacement using Phaser with the holo structure of OASS from *Brucella abortus* (PDB:5JIS) used as the search model (Dharavath et al., 2017; McCoy et al., 2007). The subsequent 1.46 Å LigB-bound and 1.60 Å acetate-bound structures were solved using the 1.47 Å LigA/holo structure for molecular replacement in Phaser. These models were subjected to multiple rounds of rebuilding in Coot and refinement using Phenix until the refinements converged (Adams et al., 2010; Emsley & Cowtan, 2004). Data collection, processing and refinement statistics are presented in Table S1.

### Cysteine Ninhydrin OASS Activity Assay

The kinetic parameters of AfuCys1a catalysing the OASS reaction were measured using the acid-ninhydrin method adapted from the methods developed by Gaitonde, 1967 and Pederick et al., 2024. All reactions were performed at room temperature (approximately 23°C). Composition of reaction mixes and preparation of solutions are detailed in the materials table. Solutions were mixed by pipetting upon addition of each reagent. Aliquots of 10 µM AfuCys1a were thawed at room temperature and diluted to 125 nM in a 2× protein reaction mix. This was transferred to PCR strip tubes in 25 µL aliquots and reactions were initiated by the addition of 25 µL of 2× substrate solution. The final 1x 50 µL reaction mixture comprised 62.5 nM of AfuCys1a, 20 mM Tris-HCl pH 7.5, 50 µM PLP, 1 mM sodium sulphide, and 0.4–25 mM L-OAS.

The reactions were left to proceed for 7 min at room temperature and then quenched simultaneously by adding 50 µL of glacial acetic acid. For colouration, 50 µL of ninhydrin acid reagent was added; samples were then heated to 99°C for 10 min in a pre-heated thermal cycler, resulting in a proportional colour change from pale yellow to pink, dependant on the concentration of L-cysteine. The samples were then incubated at 19°C for 5 minutes, briefly centrifuged, and 100 µL of each sample was added to 300 µL of absolute ethanol (−20°C) to stabilise the colour. For measurement, 300 µL of each sample was transferred to an individual well of a 96-well flat bottom plate. Absorbance was measured at 560 nm, and a negative control (1× reaction mix excluding L-OAS) was used for blank subtraction. The concentration of L-cysteine was determined by producing an L-cysteine calibration curve and fitting data to the linear equation. All concentrations of L-cysteine were resuspended in 50 µL of the calibration curve reaction mix. A sample that did not contain L-cysteine was used for blank subtraction. The latter steps were performed in an identical manner to the cysteine ninhydrin assay described above. All measured concentrations of L-cysteine corresponded to the concentration of the final 50 µL reaction mix. The L-cysteine standards were linear between 12.5 µM and 150 µM.

### Cysteine Ninhydrin OASS Inhibition Assay

Several small molecules were tested for inhibitory activity against AfuCys1a in the ninhydrin assay, including L-methionine, L-pipecolic acid, 2-picolinic acid, L-cystathionine, L-cycloserine, D-cycloserine, L-proline, pyroglutamic acid, 3-Methylpyridine-2-carboxylic acid, 6-Hydroxypyridine-2-carboxylic acid, and 3-picolinic acid. Candidate inhibitors were prepared as 100 mM stock solutions in Milli-Q water, adjusted to pH 7 if required, and stored at −20 °C. Final inhibition assay mixes contained 62.5 nM AfuCys1a, 20 mM Tris-HCl pH 7.5, 50 µM PLP, 1 mM sodium sulphide, 2 mM L-OAS, and serial dilutions between 0.125–100 mM of candidate inhibitor. A reaction mix without candidate inhibitors was included to calculate the relative remaining activity of AfuCys1a. A negative control (1× reaction mix excluding AfuCys1a) was included for blank subtraction. To assess if candidate inhibitors would affect absorbance readings, 50 µM L-cysteine was resuspended in the calibration curve reaction mix and assayed with each candidate inhibitor (∼0.125–100 mM). Absorbance readings were compared to a sample that did not include candidate inhibitors. Any relative effects on absorbance were subtracted from the absorbance reading given by the enzyme inhibition assay.

### Determination of Kinetic Parameters for L-OAS or AfuCys1a in the OASS reaction Activity Assay and Inhibition Assay

For determination of kinetic parameters (*K*_M app_ L-OAS and *k*_cat_) of AfuCys1a in the OASS reaction. Experiments were performed four times, with a single replicate measured for each concentration per experiment. Data were fit with the substrate inhibition equation **(1)** by non-linear regression in GraphPad Prism 9 (GraphPad Software, Inc.):

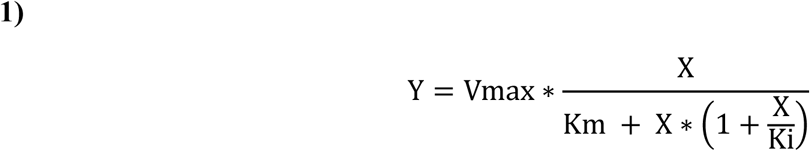

The candidate inhibitors 2-picolinic acid, 3-Methylpyridine-2-carboxylic acid, 6-Hydroxypyridine-2-carboxylic acid and 3-picolinic acid produced inhibitory effects against AfuCys1a, while the remaining molecules showed little to no inhibition. The IC_50_ value for 2-picolinic acid was determined by repeating the experiment three times with serial dilutions between 0.125–64 mM. Preliminary IC_50_ values were obtained for 3-Methylpyridine-2-carboxylic acid, 6-Hydroxypyridine-2-carboxylic acid and 3-picolinic acid, but these experiments were not repeated as inhibitory effects were not significantly more potent than 2-picolinic acid. Data were fit to the log(inhibitor) vs. response – variable slope equation **(2)** by non-linear regression in GraphPad Prism 9 (GraphPad Software, Inc.):

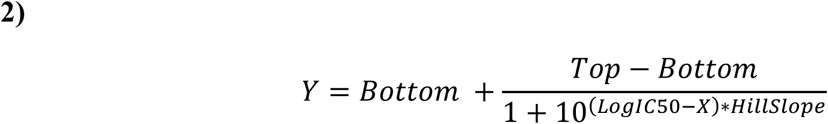

### Protein Sequence, Structure and Mechanistic Analysis

$Figures for structural visualisation were generated using PyMOL (Schrödinger, 2025). Protein secondary structures were identified using prediction by DSSP (Define Secondary Structure of Proteins) and PyMOL version 3.0 (Kabsch & Sander, 1983; Schrödinger, 2025); following identification by DSSP and PyMol, the secondary structures were then assessed by visual inspection. PubChem was used to identify candidate inhibitory molecules (S. Kim et al., 2025). Multiple sequence alignments were generated on the EMBL-EBI Web site using Clustal Omega set to default parameters (Sievers et al., 2011). Structural homologues used for structural and sequence alignments were identified by the HHpred server (Zimmermann et al., 2018). Protein sequences were retrieved from the UniProt database (The UniProt Consortium, 2023). Genomic annotations were obtained from Ensembl (Martin et al., 2023). Nucleotide and protein data were accessed via the NCBI databases (O’Leary et al., 2024). Protein structure predictions were generated using AlphaFold3 (Abramson et al., 2024). TargetP-2.0 and MULocDeep were used to predict the subcellular location of proteins based on the amino acid sequence (Armenteros et al., 2019; Jiang et al., 2023). Chemical reaction schemes were drawn using Marvin (ChemAxon) (Cherinka et al., 2019).

Molecular docking was performed using ICM-Pro (MolSoft LLC) (Abagyan et al., 1994; Abagyan & Totrov, 1994). The protein structure of the acetate-bound state was prepared by removing non-essential solvent molecules. For docking with glutathione persulphide, the PLP cofactor was replaced with the α-aminoacrylate intermediate (PDB: P1T) and PLP was removed prior to docking with the L-OSS-PLP complex. The catalytic lysine (Lys74) was positioned in the same location as structural homologues that contain a substrate in the external aldimine formation such a lanthione synthase from *Fuseobacterium nucleatum* (PDB: 5XEM) or the α-aminoacrylate in *Staphylococcus aureus* SbnA (PDB: 5D85). For ligand preparation, the SDF file of glutathione persulphide was downloaded from PubChem, which is listed as S-sulfanylglutathionate(1-), while the L-OSS-PLP complex was prepared in Marvin (ChemAxon), and saved as an SDF file. The protein and ligands were converted in ICM-Pro; hydrogen atoms were added, protonation states were assigned, and formal charges predicted in ligands. The docking search space was defined around the active site based on the position of PLP and ligands within the structures of AfuCys1a. Docking calculations were performed using a docking effort of 2 and the ICM-Pro default parameters unless otherwise stated. Generated poses were ranked according to the ICM docking score, and the highest-scoring poses with catalytically relevant binding orientations were selected for analysis and visualisation.

### Intact and Native Mass Spectrometry

For both intact mass spectrometry and native mass spectrometry, AfuCys1a was buffer exchanged into 200 mM ammonium acetate (pH 6.8) by five rounds of dilution to 15 mL followed by concentration to 0.5 mL using a 10 kDa MWCO centrifugal filter. Intact protein mass measurements were carried out under denaturing conditions using an Agilent 6230 time-of-flight instrument coupled to an Agilent 1260 Infinity II LC System. Protein sample (1 µL) was injected and electrosprayed using 50 % aqueous acetonitrile/0.01 % formic acid at a flow rate of 0.2 mL/min, without chromatographic separation. ESI-MS conditions were: positive-ion mode; capillary voltage, 3500 V; nozzle voltage 2000 V; fragmentor, 175 V; gas 8 L/min; gas temperature, 325 °C; sheath gas 11 L/min; and sheath gas temperature, 350 °C, m/z range, 500-3200. Spectra were deconvoluted using the Maximum Entropy algorithm in BioConfirm software (Agilent), over the mass range 10000-50000 Da. Native mass-spectrometry was performed on a Bruker Impact II HDMS Q-ToF mass spectrometer (Bruker Daltonics) with a nanoelectrospray ionisation source with the following instrument parameters: m/z range, 500-10,000; polarity, positive; capillary voltage, 2 kV; end plate offset, 500 V; and source temperature, 60 °C. Protein samples were electrosprayed from platinum-coated borosilicate glass capillaries (Harvard Apparatus, USA) prepared in-house. All spectra were analysed using the Compass Data Analysis software (v4.2 Bruker Daltonics). Deconvolution of native mass spectra was performed using UniDec (v6.0.3 University of Arizona) (Marty et al., 2015) with the following parameters: *m/z* range, 1,000-10,000; charge range; 1-50; sample mass every 1 Da; peak detection range 1 Da; charge smooth width, 1; peak width. 0.85.

## Results and Discussion

### Characterisation of AfuCys1a by X-ray crystallography

We identified the protein with Uniprot ID:Q4WE91 as a putative Cys1a from *A. fumigatus* (AfuCys1a). Protein expression and purification of AfuCys1a produced two distinct species, suggestive of an unexpected proteolysis event. The first species was around the expected molecular weight of AfuCys1a with N-terminal 6xHis tag at 41.8 kDa, while the second species was closer to 40 kDa and substantially more abundant. Fractions containing the ∼40 kDa species were pooled and used for all subsequent experiments discussed herein. Intact mass spectrometry (MS) was performed to determine the exact molecular weight of the ∼40 kDa protein, revealing these fractions comprised two further proteoforms of AfuCys1a at 39.150 and 39.298 kDa. These masses indicate the occurrence of a spontaneous cleavage event at the sites Arg8/Phe9 and Phe9/Gly10 during expression and purification, as well as the pyridoxal-5′-phosphate (PLP) cofactor induced covalent modification of AfuCys1a (Figure 1). Analysis of an AlphaFold3 model, combined with signal sequence predictive server TargetP 2.0, and protein localisation prediction tool MULocDeep, suggested that a mitochondrial target sequence (MTS) is present at the N-terminal region of AfuCys1a, and this enzyme localises to the mitochondrial matrix. Cleavage or proteolysis events are often favoured in disordered or highly flexible regions such as MTSs, and would reasonably explain the loss of the first 8–9 N-terminal residues during protein expression and purification (Francis & Page, 2010; Gottesman, 1990).

**Figure 1:**
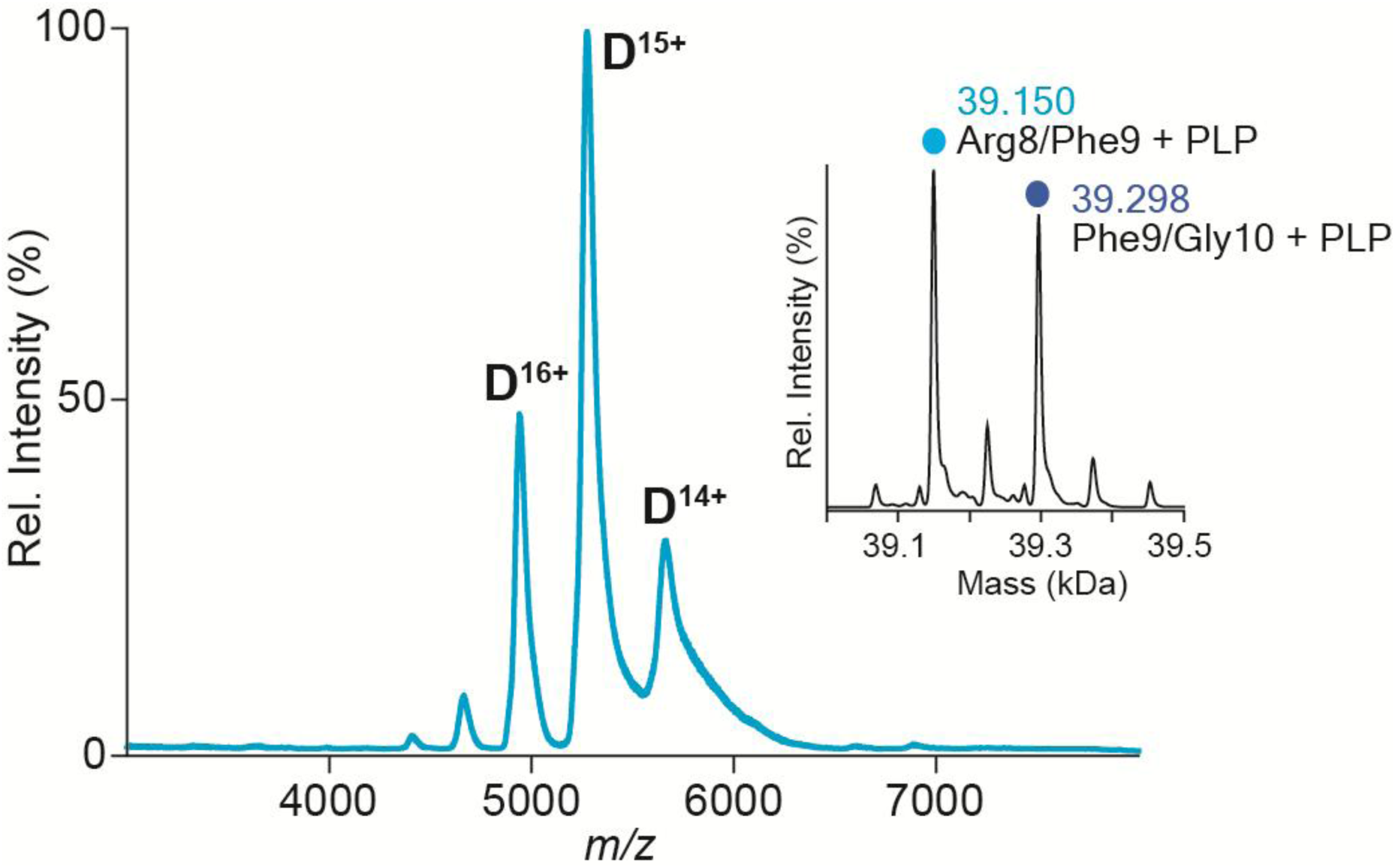
Quaternary State of AfuCys1a is Exclusively Dimeric. Native MS shows AfuCys1a (20 μM in 200 mM ammonium acetate, pH 6.8) oligomeric state as a folded dimer (blue). **Inset:** Intact MS and mass deconvolution of PLP modified AfuCys1a shows the spontaneous cleavage at Arg8/Phe9 (light blue) and Phe9/Gly10 (dark blue).

For structural characterisation of AfuCys1a, X-ray crystallography was conducted, yielding structures of four distinct ligand-bound states derived from three high-resolution dimeric structures (1.46–1.79 Å). The cofactor pyridoxal-5′-phosphate was covalently bound in all monomers, each crystal contained two polypeptide chains in the asymmetric unit, and all structures were solved as antiparallel dimers. The dimeric formation of AfuCys1a was experimentally validated by native MS which indicated that AfuCys1a is predominantly present as dimer bound with PLP in solution with a molecular weight of ∼79.2 kDa (Figure 1).

For these experiments, the original aim was to obtain the holo structure of AfuCys1a, where the protein is linked to its cofactor PLP only, with no other ligands present in the active site. Instead, distinct molecules were present in the active site that either co-purified with AfuCys1a or were present in the crystallisation condition. Four distinct ligand bound states are discussed here: acetate-bound, LigA-bound, LigB-bound, and holo. Crystallographic data collection, processing and refinement statistics for each structure are presented in Table 1.

**Table 1.**
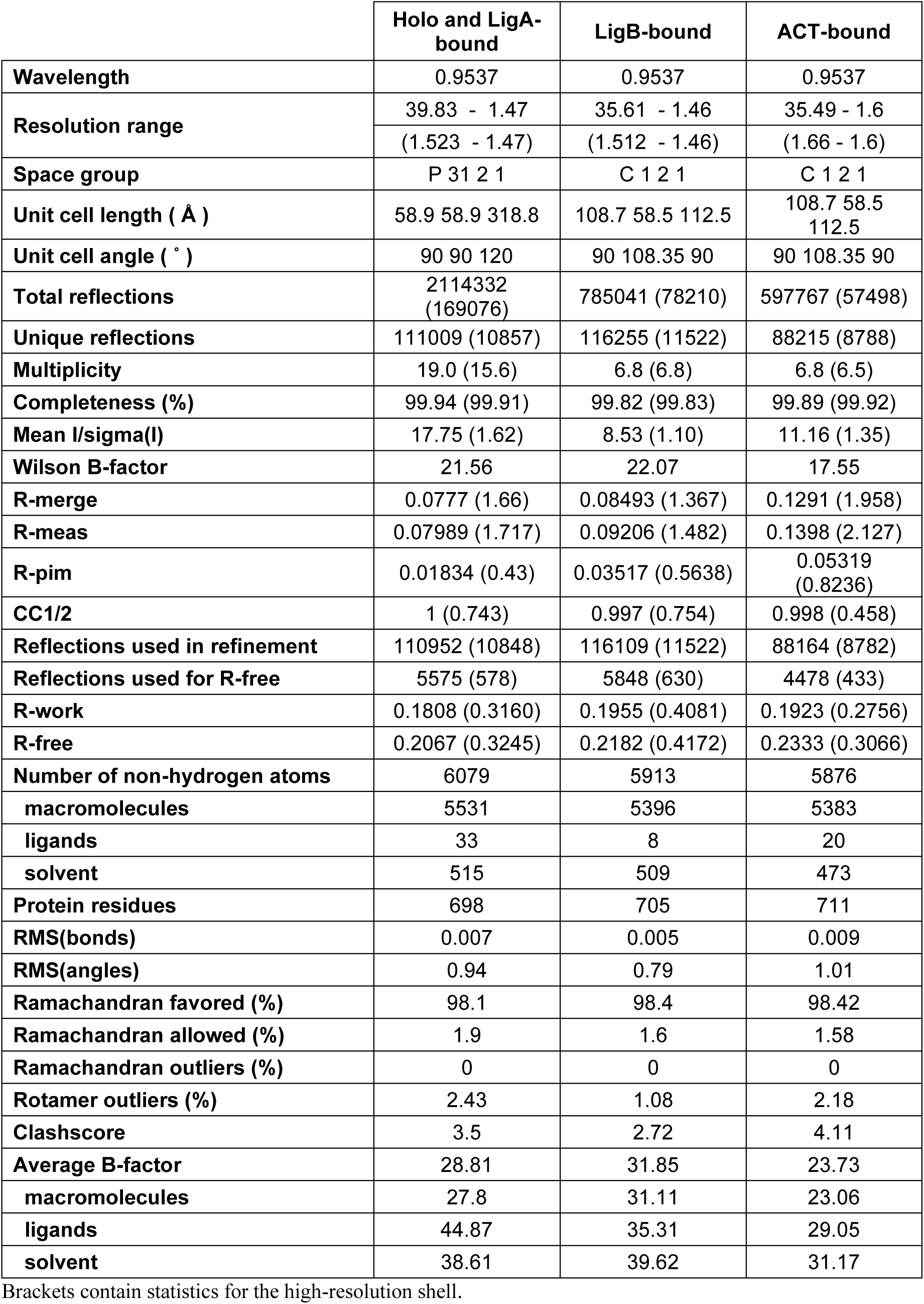
Data collection and Refinement Statistics for the Holo and LigA-bound Structures, the LigB-bound Structure, and the ACT-bound Structure.

LigA and LigB represent an unidentified cyclic molecule, which is likely the same cyclic compound. In total, this compound was observed in the active site of five monomers of AfuCys1a, obtained from three different crystallisation conditions. Analysis of the *2F_o_-F_c_* and *F_o_-F_c_* electron density maps suggests that LigA and LigB form a 5-membered ring and comprise a carboxyl group. No molecule matching this structure or these assumed characteristics were present among any of the reagents used during protein purification or crystallisation, suggesting that LigA and LigB may have originated from *E. coli* during protein expression. Although this narrows the list of possible identities, several *E. coli*-derived molecules could still account for the observed electron density. The cyclic compound could not be confidently identified, and consequently, the two molecules discussed here are referred to simply as LigA and LigB, to distinguish them within their respective protein structures.

The acetate-bound state was solved to a resolution of 1.6 Å, and contained PLP, a sodium ion (Na^+^) and an acetate molecule within the active site. The LigA-bound and holo states are form the same dimeric structure which was solved to a resolution of 1.47 Å. The LigA-bound state contains PLP and the unidentified cyclic molecule (LigA) within the active site, while the holo state contains PLP and a potassium ion (K^+^) within the active site. The holo state also contains a glycerol molecule which is located in close proximity to the catalytic site. Lastly, the LigB-bound state was solved to a resolution of 1.46 Å and contained PLP, Na^+^ and an unidentified cyclic molecule (LigB).

The structure of AfuCys1a displays an overall fold characteristic of PLP-dependent fold type II or Tryptophan-β-synthase family proteins (Eliot & Kirsch, 2004; Liang et al., 2019; Schneider et al., 2000). The acetate-bound state is used here for the assignment of secondary structures and is representative of all structures discussed here (Figure 2). Each monomer folds into two sub-domains, sub-domain 1 and sub-domain 2, which each consist of an α/β domain. The secondary structures comprise a repeated α-β-α-β motif until the C-terminal region which shows the organisation of α11-β10-α12-α13 (Ala308–Pro371). The active site is formed by multiple secondary structures from both sub-domain 1 and 2, which forms a cleft-like structure, encompassing the active site. Within sub-domain 1, the first residue of helix 3 is the catalytic lysine, Lys74, which forms the PLP internal aldimine; for helix 4, first residue is Thr105 which is part of the Asparagine Loop TAGNT (residues 101–105), a highly conserved substrate loop within PLP-dependent fold type II family proteins. Within sub-domain 2, the last two residues of the PLP phosphate binding site GTGGT (residues 210–214) form the start of helix 8, and the second substrate loop SITEGIG (residues 259–265) is located within a flexible region (Gly249–Asp281) between helix 9 and beta-strand 9.

**Figure 2.**
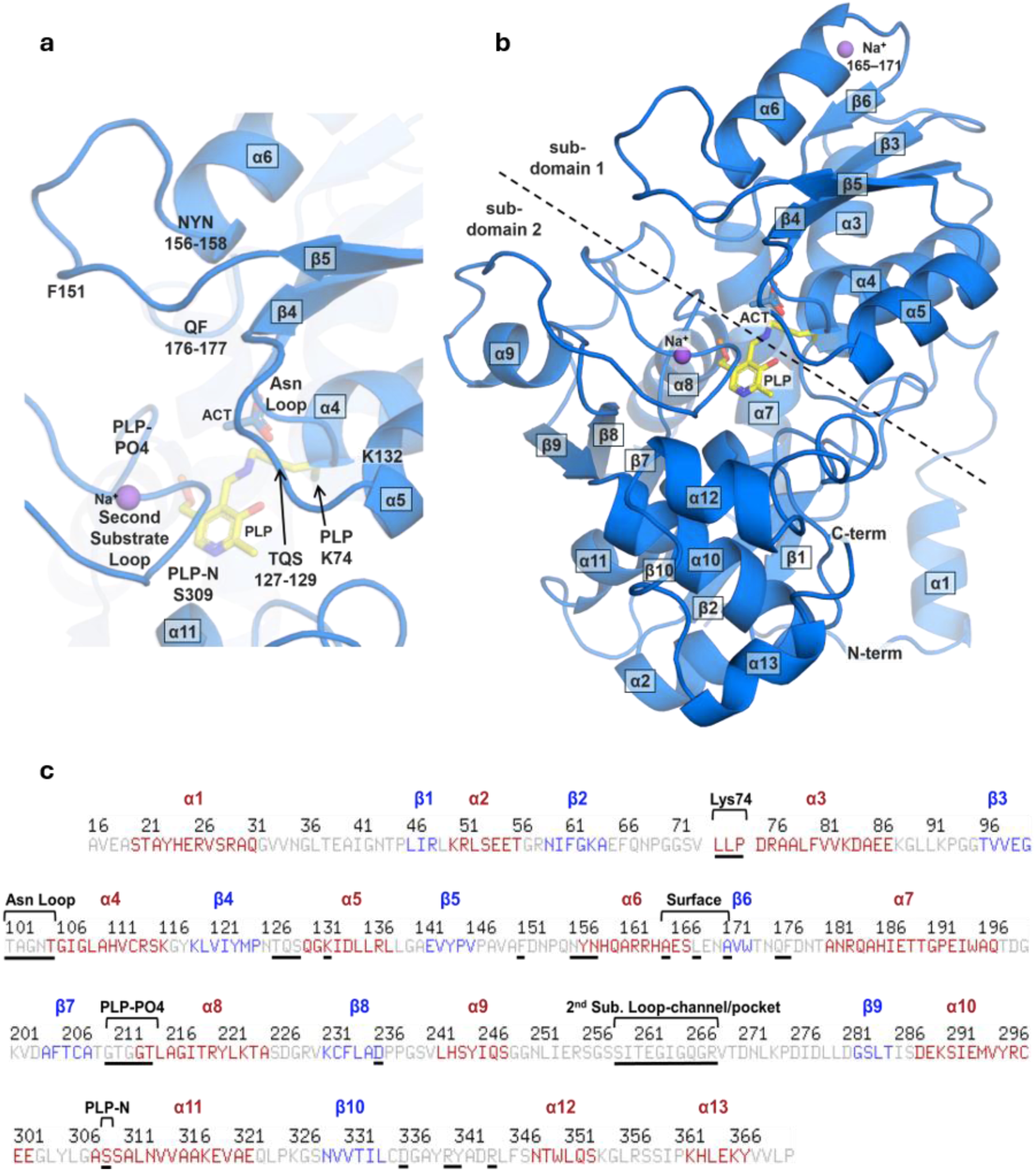
The Overall Fold of AfuCys1a and Location of Key Features. **a** Close-up of the active site of the acetate-bound state of AfuCys1a. Key features, residues and ligands are labelled. **b** Assigned alpha helices of the acetate-bound state of AfuCys1a are numbered from the N-terminus to the C-terminus and labelled α1–α13, while beta-strands are labelled β1– β10. **c** Residues of the AfuCys1a amino acid sequence that form alpha helices are coloured red, while residues that form beta-strands are coloured blue. Active site residues are underlined in black and features are labelled.

Several ligand binding sites are identified in the structures of AfuCys1a due to the presence of distinct molecules within the protein structures. One such molecule is the cofactor PLP, which is covalently bound in all structures of AfuCys1a by formation of an internal aldimine between the ε-amino group of Lys74 and the aldehyde group of PLP. This cofactor was intentionally supplemented throughout protein expression and purification. Several active site residues are involved in PLP binding: the side chain of Asn104 (within the Asparagine Loop) forms a hydrogen bond with the PLP phenolic 3-hydroxyl group, the residues GTGGT (residues 210–214) anchor the PLP phosphate group, and the sidechain of Ser309 forms a hydrogen bond with the N of the PLP pyridine ring.

For PLP-dependent fold type II enzymes, the Asparagine Loop is known to bind its first substrate by forming hydrogen bonds with an amino acid carboxyl group (Burkhard et al., 1999; Devi et al., 2019; Feldman-Salit et al., 2009; Mazumder & Gourinath, 2016; Meier et al., 2001; Tran & Brown, 2022). Placement of acetate in the acetate-bound state is supported by its presence in the structures of multiple homologues (PDB: 6C2Z; 6VJU; 4COO; 1SFT). The carboxyl groups of acetate, LigA and LigB can participate in hydrogen bonds with the Asparagine Loop through the backbone amides of Ala102, Asn104 and Thr105, the side chain of Thr101. The ligand carboxyl groups are also stabilised by hydrogen bond formation with the side chain of Gln176, located adjacent to the Asparagine Loop. Additionally, the sidechain of Gln176 also forms a hydrogen bond with the side chain of Thr105, suggesting that Gln176 may contribute to structural stability of the Asparagine Loop. For carboxyl-binding substrate loops, a similar mechanism of hydrogen bond formation has been described previously, where crowded amide-carboxyl interactions form a flexible adaptive hydrogen bonding network with different atoms interchanging between donors and acceptors (Kurkcuoglu et al., 2012; Romero-Rivera et al., 2022).

Metal coordination with either K^+^ or Na^+^ was identified at two separate sites over several structures of AfuCys1a. The first was a protein surface metal binding site, formed by the backbone carbonyls of Ala165, Leu168 and Ala171. The second metal binding site is within the second substrate loop and formed through the backbone carbonyls of Ser259 and Thr26. For the LigA-bound and holo state, K^+^ is modelled and supported by peaks in the 2 *F_o_-F_c_* and *F_o_-F_c_* electron density map which are 2–3 times stronger than nearby water molecules, consistent with the relative masses between potassium and water (S. Kim et al., 2025; Terwilliger & Berendzen, 1999). Furthermore, most K^+^ ions were observed to have either octahedral or square pyramidal coordination geometry with distances measured between 2.6–3 Å, characteristic of K^+^ coordination bonding (Harding, 2002, 2006; Vadivel et al., 2019). For the LigB-bound and acetate-bound states, Na^+^ was modelled at these sites as sodium was present in buffers and crystallisation conditions. Relative strength in the electron density maps could not be used to support placement of sodium as their masses are near identical. Instead, placement of these ions were supported by close proximity with 2 or 3 backbone carbonyls as well as nearby water molecules at distances consistent with ionic bonding for Na^+^, between 2.25–2.7 Å (Harding, 2002, 2006). A metal ion was present within the second substrate loop of all states of AfuCys1a except the LigA-bound state.

Lastly, a glycerol molecule was identified in the holo structure of AfuCys1a in a pocket located adjacent to the active site substrate loops. The glycerol molecule forms hydrogen bonds with the side chains of Arg268 and Gln266 as well as engaging in van der Waals interactions with the side chains of Phe177 and Phe151.

Within the active site, several residues are present in identical conformations between AfuCys1a states (Figure 3): Phe151, Asn156, Tyr157, Asn158, Gln176, Phe177, Tyr341 and Asp336. The consistency of these residues could suggest a role in maintaining overall structural stability or specific catalytic functions where precise positioning is crucial (Yabukarski et al., 2020; Yang & Bahar, 2005). In comparison, the residues: Thr127, Gln128, Ser129, Lys132, Phe177, Arg340, Tyr341, and Arg344, were identified in alternative conformations between AfuCys1a states. Flexibility of these residues could be associated with substrate binding and catalysis, where mobility allows adaptation to different substrates, transition states, or products (Johnson, 2008; Riziotis et al., 2022; Yabukarski et al., 2020; Yang & Bahar, 2005). Similar flexibility and conformational changes are demonstrated by the Asparagine Loop, the second substrate loop and residues that comprise the channel/pocket (Figure 3). Flexibility is a common feature of substrate loops and is often closely linked to substrate recognition, catalysis, and dynamic regulation of enzyme function (Gunasekaran et al., 2003; Gunasekaran & Nussinov, 2004; Malabanan et al., 2010). Ultimately, a balance between stable and flexible residues is essential for both enzyme function and overall structure. These general associations between structure and function are not definitive by any means but can be a valuable starting point for characterisation of novel enzymes such as AfuCys1a (Hou et al., 2023; Johnson, 2008).

**Figure 3.**
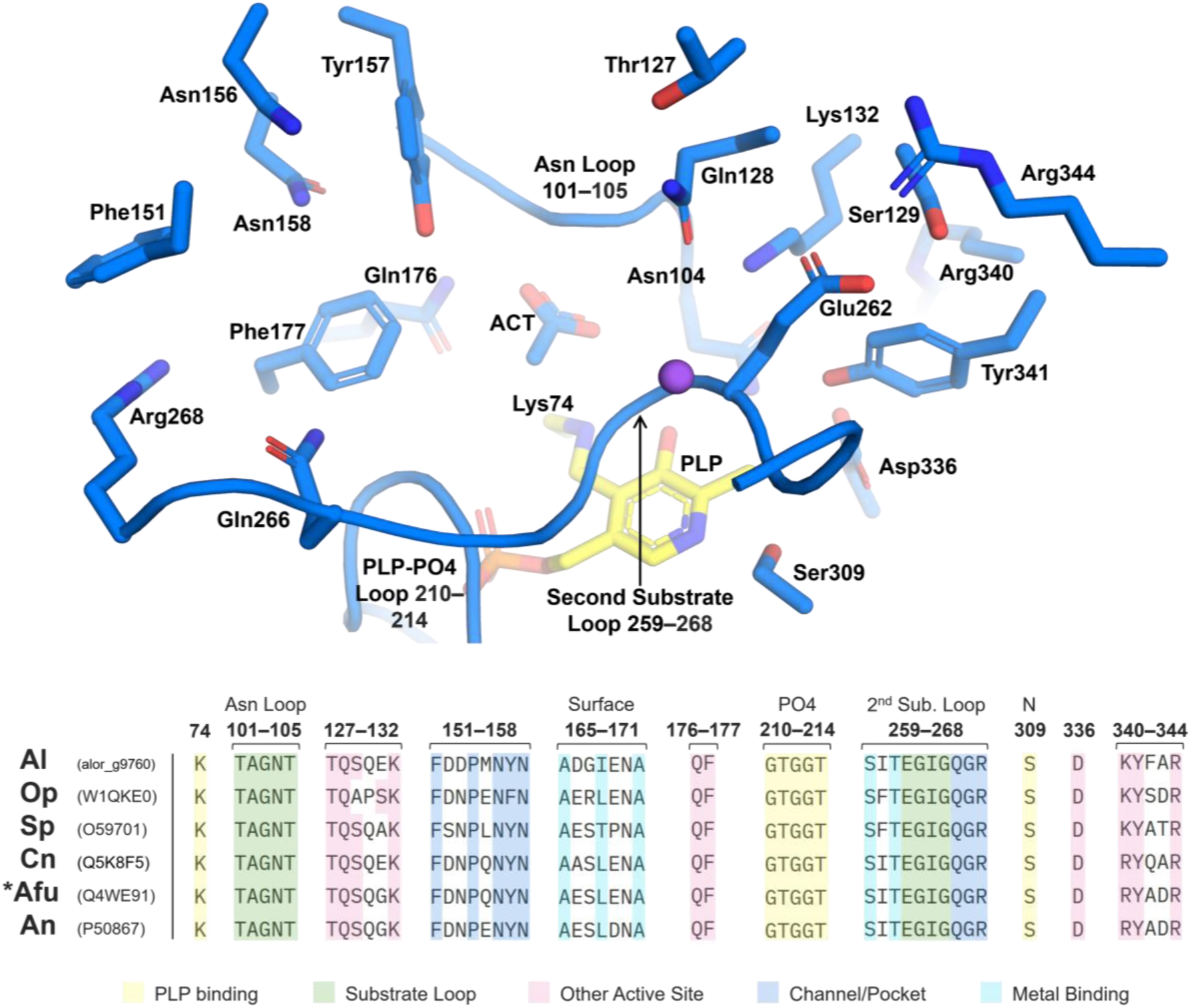
Spatial Positioning of Active Site Residues in the Acetate-bound state of AfuCys1a and Multiple Sequence Alignment of Fungal Cys1a Enzymes. Amino acid side chains are individually labelled, and regions are labelled for the Asparagine Loop, the second substrate loop and the PLP phosphate binding loop. Uniprot codes are provided next to each Cy1a enzyme included in the multiple sequence alignment.

Key insights into structure-function relationships can be inferred by aligning the sequence and structure of AfuCys1a with those of close homologs from other species (Borkakoti et al., 2025). To date, partial characterisation has been achieved for three Cys1a enzymes: AfuCys1a, SpCys1a from *Schizosaccharomyces pombe*, and lastly, ACys1a from the coral *Acropora loripes* (Bastard et al., 2017; Salazar et al., 2022). By aligning the Cys1a sequences and integrating available functional data with structural information from AfuCys1a, important insights can be gained (Figure 3). Such analyses can reveal conserved residues that are potentially crucial for catalytic activity, support the characterisation of additional Cys1a enzymes through sequence homology, and identify sequence variations that could indicate species-specific adaptations or functional divergence (Marini et al., 2010; Stamboulian et al., 2020).

Sequence alignment included the partially characterised Cys1a enzymes, AfuCys1a, SpCys1a and ACys1a. For the purpose of further characterising enzymes of this class, putative fungal Cys1a enzymes were also included in the sequence alignment (Figure 3). These putative fungal Cys1a enzymes were previously broadly classified cysteine synthases and include Cys1 from *Ogataea parapolymorpha* (OpCys1), Cys1 from *Cryptococcus neoformans* (CnCys1), and CysB from *Aspergillus nidulans* (AnCysB) (Bastard et al., 2017; Borkovich et al., 2004; Brzywczy et al., 2007; Radford, 2004; Salazar et al., 2022; Sievers et al., 2011; Sohn et al., 2014; Toh et al., 2023). Alignment indicated that these enzymes shared a minimum of 52 % sequence identity and near identical conservation of active site residues. Results of this comparative sequence alignment in combination with high-resolution X-ray crystal structures support the identification of OpCys1, CnCys1 and AnCysB, as Cys1a enzymes, and will herein be referred to as OpCys1a, CnCys1a and AnCys1a, respectively.

### LigA and LigB Cause Disruption While Metal Binding Stabilises the Active Site

Alignment of AfuCys1a ligand-bound states allowed identification of conformational changes associated with specific ligand binding. The second substrate loop of AfuCys1a was compared between the ligand bound states. For the acetate-bound and holo structure, the second substrate loop is positioned close to the centre of the active site. For LigA- and LigB-bound structures, the second substrate loop appears to be displaced by LigA and LigB; if the second substrate loop were present in the same conformation as the holo and ACT-bound states, LigA and LigB would clash with the backbone carbonyl of Gly263 (within the second substrate loop) at 1.8–2.1 Å, leading to a shift in its conformation away from the centre of the active site (Figure 4a). However, this disruption may be partially rescued by metal binding within the second substrate loop. This is known function of metal coordination where ions such as K⁺ or Na⁺ help preserve protein structure, stabilise substrate-binding regions, and sometimes participate directly in catalysis by coordinating with backbone carbonyls or water molecules (Dutta & Bahar, 2010; Page & Di Cera, 2006; Permyakov, 2021; Vadivel et al., 2019; Woehl & Dunn, 1995). The impact of metal binding at the second substrate loop can be demonstrated by comparison between the LigA- and LigB-bound states. In both states, the unidentified cyclic molecule displaces the second substrate loop when compared to the holo or ACT-bound structures. However, this displacement is less pronounced in the LigB-bound state, which also binds Na^+^ at the second substrate loop. In contrast, no metal ion is present at the second substrate loop within the LigA-bound state, indicating that enhanced stability is afforded by metal binding at this site.

**Figure 4.**
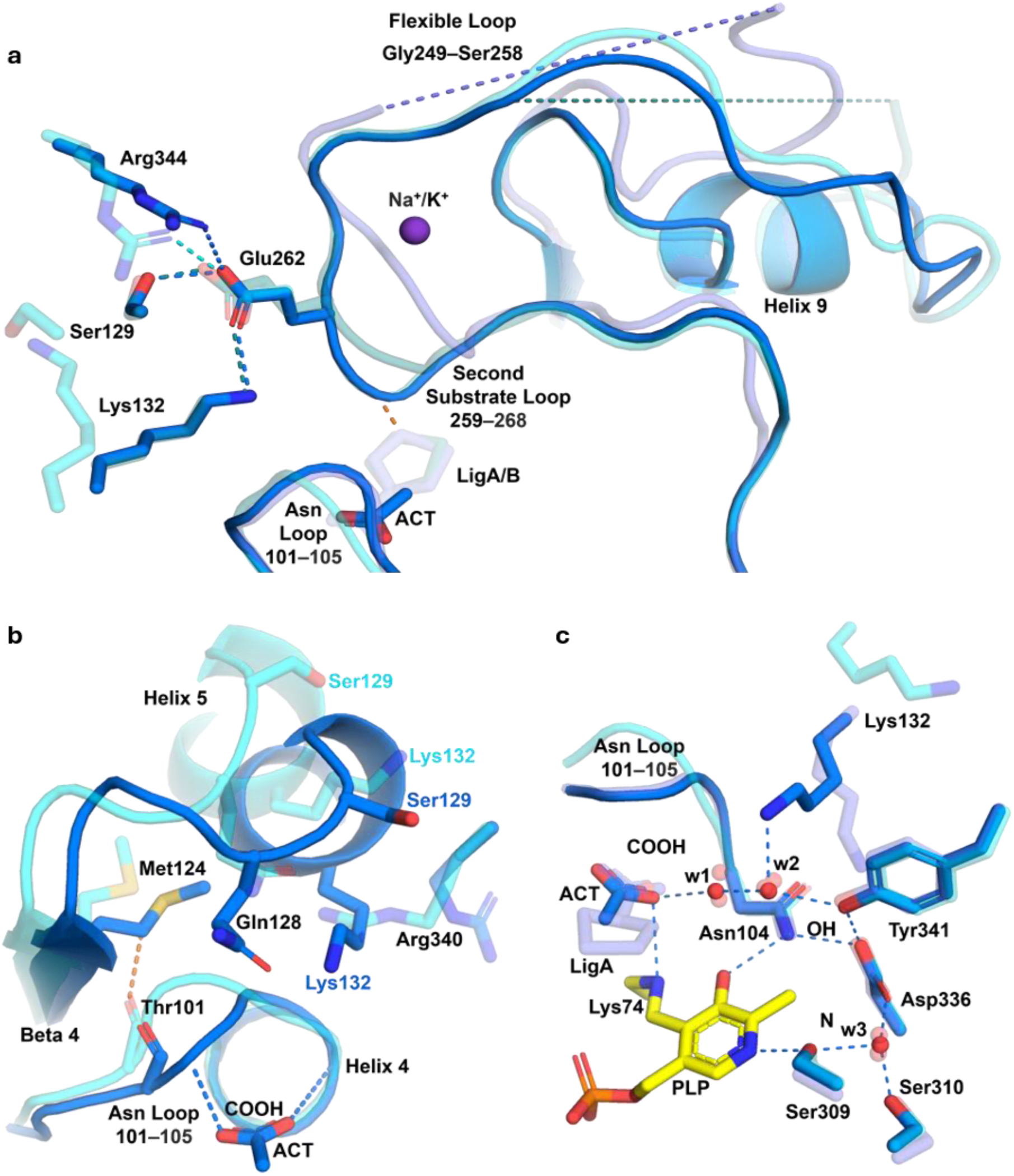
Superimposition of ligand Induced Conformational Change of AfuCys1a. The acetate-bound state is coloured dark blue. The three other states are transparent and include the holo state (cyan), the LigA-bound (slate) and the LigB-bound (teal). Hydrogen bonds are dashed lines in blue, and steric clashes are dashed lines in orange. **a** Variable hydrogen bond formation of Glu262 and metal binding at the second substrate loop is associated with increased stability of the second substrate loop, the flexible loop (Gly249–Ser258) and helix 9, while LigA and LigB are associated with disruption. **b** Movement of helix 5 and the Asparagine Loop away from the active site is associated with absence of a ligand carboxyl group (COOH) and Thr101 clashing with Met124. **c** Residues and conserved positions of water molecules (w1–w3) that form an extensive hydrogen bonding network involving the Asparagine Loop, the carboxyl group of acetate (ACT), the Lys74 ε-amino group, and the nitrogen (N) of the PLP pyridine ring.

Ligand binding to the second substrate loop is also linked to larger conformational changes, impacting regions such as Gly249–Ser258 which precedes the second substrate loop, as well as helix 9. Alignment of the four AfuCys1a states indicated that Glu262 within the second substrate loop is a key residue for stability (Figure 4a). For the LigB-bound and ACT-bound state, the side chain of Glu262 forms a hydrogen bond with the side chain of Lys132 or with Ser129 (within helix 5 and the preceding loop). For the holo state, AfuCys1a adopts an ‘open’ conformation that positions Lys132 and Ser129 outside of the active site; instead, the side chain of Glu262 forms a hydrogen bond with Arg344. In contrast, the side chain of Glu262 could not be modelled for the LigA-bound state. Despite this, it can be inferred from the backbone position that Glu262 would not be able to participate in hydrogen bond formation in the LigA-bound state. The absence of Glu262 hydrogen bond formation in the LigA-bound state is associated with disruption to the second substrate loop, loss of metal binding, and increased disorder within the region Gly249–Ser258. This may also suggest that reduced displacement of the second substrate loop in the LigB-bound state is linked to both metal binding and Glu262 hydrogen bond formation, which also stabilise one another. Decreased stability of the second substrate loop and the region Gly249–Ser258 in the LigA-bound state is further exemplified by disruption of helix 9, which could not be modelled from Ile246 to Ser248, and did not adopt a secondary structure. However, in the LigA-bound state, crystal contacts are present at the location of helix 9, forming hydrogen bonds with the side chains of Tyr245 and His243. Although helix 9 undergoes substantial shift linked to ligand binding in the active site, the loss of secondary structure is likely due to crystal contacts in this area rather than intrinsic instability of the helix.

Ligand induced conformational changes were also observed for the Asparagine Loop. In comparison to the holo state, binding of a ligand carboxyl group (acetate, LigA, and LigB) to the Asparagine Loop stabilises the loop a ‘downward’ conformation, positioning it closer to PLP and the centre of the active site. This conformational switch is widely conserved among this structural class (PDB: 8SRU; 6C2Z; 5D85; 2Q3D; 3VSD; 2V03; 4COO; 8B9M), indicating that carboxyl group binding to the Asparagine Loop plays broad functional role in the binding and positioning of distinct ligands. The holo state of AfuCys1a shows that when a ligand is not bound, the Asparagine Loop is no longer maintained closer to the active site, and this is associated with a conformational change of helix 5 away from the active site. Superimposition demonstrated that the backbone carbonyl of Thr101 (within the Asparagine Loop) of the holo state would clash with the side chain position of Met124 in the other states of AfuCys1a (Figure 4b). This results in displacement of Met124 and causes subsequent steric clashes with surrounding residues including Pro145 (beta-strand 5) and Ile133 (helix 5). This then causes a knock-on effect, resulting in further clashing with residues that are either part of or near helix 5, consequently pushing helix 5 away from the active site. Movement of helix 5 causes repositioning of several active site residues away from the catalytic centre, including Thr127, Gln128 and Ser129, which comprise the loop immediately preceding helix 5, as well as Lys132 which is part of helix 5. Repositioning of these residues then allows Arg340 to occupy the newly available space.

Ligand binding and conformation of the Asparagine Loop may also be impacted by formation of a conserved proton transfer network (Figure 4c). The Asparagine Loop and the ligand carboxyl group are part of a larger hydrogen bonding network which includes the hydrophilic side chains Ser310, Asp336 and Tyr341, three water molecule positions (w1, w2 and w3), and transiently with Lys132 (within helix 5). Except for Lys132 (which is subject to ligand induced conformational change), the conformation of this network is conserved in all four states of AfuCys1a, including the holo state where the ligand carboxyl group is replaced by two water molecules. For the acetate-bound state, extensive hydrogen bond formation was observed. Here, w2 forms three strong hydrogen bonds, one with w1 at 2.4 Å, another with the side chain of Tyr341 at 2.8 Å, and lastly with the side chain of Lys132 at 2.7 Å.

Orientation and positioning of each ligand (acetate, LigA, and LigB) was also compared between the states of AfuCys1a, indicating ligand specific variation in preferred hydrogen bond formation with backbone amides of Asn104 and Thr105 (within the Asparagine Loop), the water molecule at position w1, and the N of the Lys74 imine bond with PLP. This suggests that water molecules at conserved positions (w1 and w2) may also impact ligand-specific orientation in the context of a conserved hydrogen bonding network (Figure 4c). Dense hydrogen bonding networks such as this, which include short hydrogen bond distances between charged residues and water molecules, are characteristic of proton transfer networks (J. K. Kim et al., 2025; Luang et al., 2025; Zhao et al., 2005). This would be consistent with the function of PLP-dependent enzymes, where proton transfer is crucial during the catalytic cycle for substrate positioning, stabilising intermediates and facilitating transformations such as transamination or β-replacement (Li & Sun, 2024; Mueser et al., 2020; Rabeh et al., 2005). The structures of AfuCys1a were also aligned with homologous (PDB: 8SRU; 1OAS; 2V03; 4COO), demonstrating conservation of analogous hydrogen bonding networks between related enzymes (Burkhard et al., 1998; Pederick et al., 2024; Zocher et al., 2007). In particular, the residue at AfuCys1a position Lys132 is also moderately conserved between these homologues as either a lysine or arginine, and similarly flexible and transient within the network. Collectively, structural analysis of the four states of AfuCys1a, together with the observed conservation among homologues, suggests that this network may have functional relevance within the active site of PLP-dependent fold type II enzymes.

Taken together, ligand binding to the Asparagine Loop stabilises AfuCys1a in a ‘closed’ state, adopting a position that narrows the active site cleft and properly aligns key catalytic residues within the active site. This is supported by comparison with several homologues, including mammalian serine racemase, *Entamoeba histolytica* OASS, *S. aureus* SbnA, and *Mycobacterium tuberculosis* CysM, which also adopt a ‘closed’ conformation upon ligand binding (Ågren et al., 2009; Brunner et al., 2016; Goto et al., 2009; Kobylarz et al., 2016; Raj et al., 2013; Smith et al., 2010). Additionally, helix 5 could undergo a more substantial ligand-induced conformational change than is observed in the holo-state. Residues close to or within helix 5 appear to be influenced by symmetry coordinates within the crystal lattice, which include Ser248, Asn251, Leu252 and Glu254 (comprising part of the flexible loop Gly249–Ser258). One hydrogen bond is formed between the side chain of Glu254 (symmetry coordinate) with the backbone amide of Gln130 (within helix 5) at 2.8 Å. Also, several prospective steric clashes could occur between symmetry coordinates (Ser248–Glu254) with residues between Asn126–Ser129, indicated by distances of 3.0–3.3 Å.

This comparison suggests that LigA and LigB cause disruption to the active site architecture which is partially rescued by the stabilising effect of metal binding and Glu262 hydrogen bond formation. Interestingly, no analogous ligand-induced disruption of the second substrate loop could be identified in structural homologues when bound to biologically relevant substrates. Comparable disruption to the second substrate loop was only identified in the homologue *Mt*CysM (PDB: 5I6D), which contains a designed inhibitor (compound 5) rather than a natural substrate (Brunner et al., 2016). Collectively, this suggests the unidentified cyclic molecule may act as an inhibitor against AfuCys1a and could provide a structural basis for further development.

### PLP-dependent Reaction Mechanism of AfuCys1a

The PLP cofactor is widely conserved within this structural family and necessary for catalysis. The reaction mechanism suggested here is based on the OASS ping-pong β-replacement mechanism which uses L-OAS as the first substrate and H_2_S as the second substrate (Burkhard et al., 1998; Schnell et al., 2007). For the AfuCys1a reaction mechanism, L-OSS is presented as the first substrate and deprotonated glutathione persulphide (GSS^-^) as the second substrate. The chemistry of each step in a β-replacement reaction for this family of proteins is built around the same general mechanistic framework but with specific variations for individual enzyme and its physiological substrates (Schnell et al., 2015).

As seen in the distinct states of AfuCys1a, the reaction starts with PLP in the form of an internal aldimine with the catalytic lysine (Lys74). Upon entering the active site, the α-amino group of OSS is deprotonated by an active site base (Figure 5, reaction step 1).

**Figure 5.**
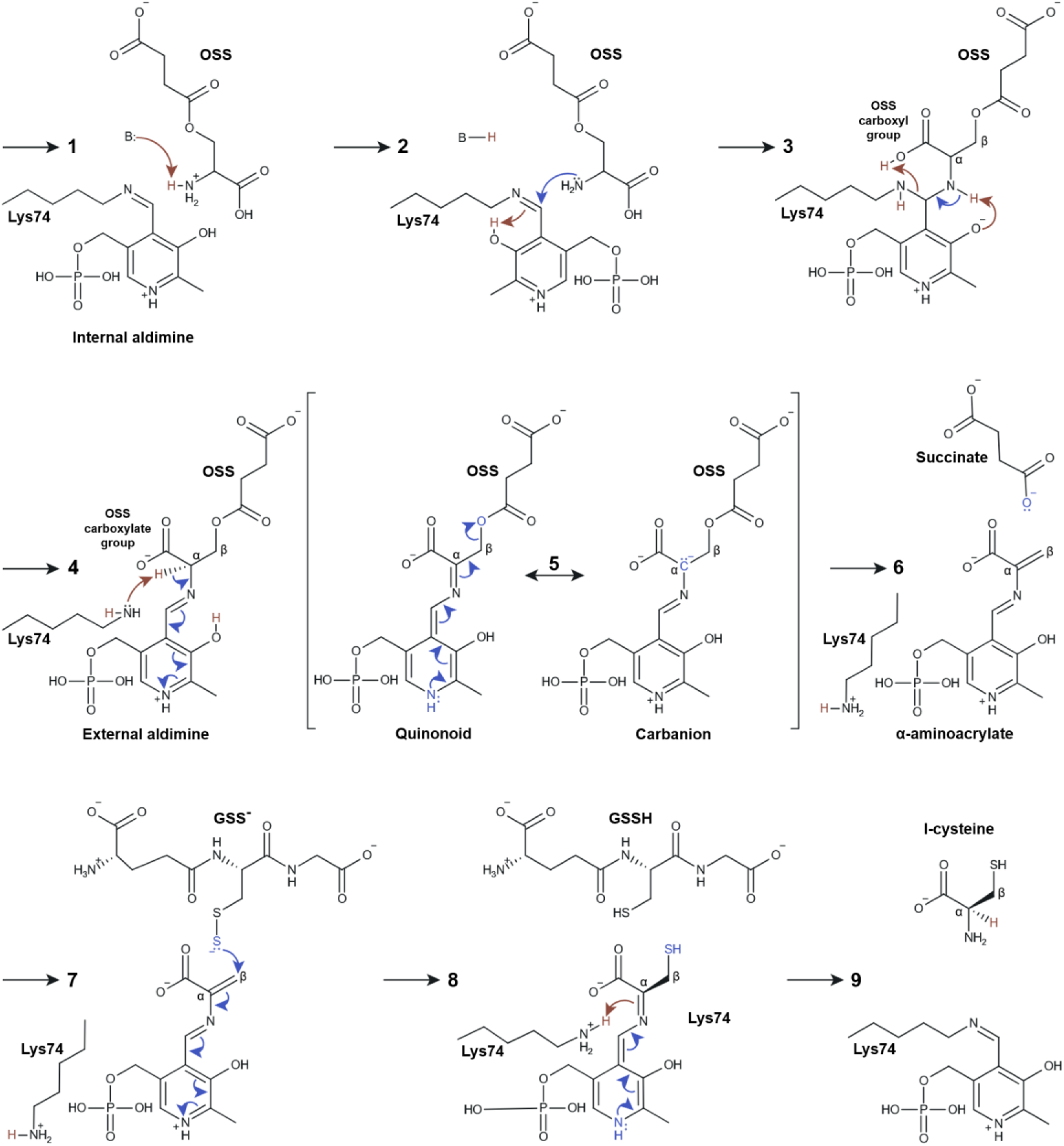
Proposed Reaction Mechanism of AfuCys1a. Curved arrows and atoms are coloured blue to represent electron transfer and coloured red when proton (H^+^) transfer is also taking place.

The deprotonated α-amino group of OSS can now perform nucleophilic attack on the Schiff base (imine) bond between PLP and the ε-amino group of Lys74. This allows the Lys74 ε-amino group to accept a proton (H^+^) from the PLP phenolic 3-hydroxyl group, deprotonating the PLP group to 3-phenolate. (Figure 5, reaction step 2).

The α-amino group of OSS is then deprotonated a second time by the PLP 3-phenolate group, resulting in electron transfer to the Lys74 ε-amino group. The Lys74 ε-amino group then accepts a proton (H^+^); it is suggested here that proton donation could occur through the ligand carboxyl group (COOH) which orientates toward the Lys74-PLP imine bond in the ligand bound structures of AfuCys1a and participates within a dense hydrogen bonding network (Figure 5, reaction step 3).

The Lys74-PLP imine bond is now broken, and the external aldimine is formed between OSS and PLP. The Lys74 ε-amino group performs nucleophilic attack on the α-carbon of OSS, accepting the α-carbon proton (H^+^) and leaving as NH^3+^. The Lys74 nucleophilic attack results in electron transfer from the OSS α-carbon to the nitrogen within the PLP pyridine ring (Figure 5, reaction step 4).

A resonance state between quinonoid and carbanion intermediate is formed by electron transfer between the OSS α-carbon and the nitrogen within the PLP pyridine ring (Figure 5, reaction step 5).

The carbanion like aromatic state of the PLP ring is more stable, favouring electron transfer to the OSS β-carbon. Succinate is a good leaving group and exits the active site with a long pair of electrons, resulting in an electron deficient β-carbon and highly reactive α-aminoacrylate intermediate that is primed for nucleophilic attack (Figure 5, reaction step 6).

Now that succinate has left, the second substrate can enter the active site. For AfuCys1a, this is likely a larger reactive sulphur species (RSS). Glutathione persulphide is used here as an example as this has been experimentally validated as a substrate of Cys1a enzymes (X. Zhang et al., 2021). Glutathione persulphide has a low p*K_a_* of 5.45 for the terminal sulphur, and is 99% present as the deprotonated form (GSS^-^) at physiological pH (Benchoam et al., 2020). The substrate GSS^-^ would readily perform nucleophilic attack on the β-carbon of the α-aminoacrylate, donating its terminal sulphur (Micheal addition). The intermediate is then stabilised by electron transfer to the N of the PLP pyridine ring (Figure 5, reaction step 7).

After donating its terminal sulphur, GSS^-^ leaves as glutathione (GSH), which is frequently protonated at its terminal sulphur at physiological pH (Benchoam et al., 2020). Electrons transfer back to the α-carbon of the intermediate which can then accept a proton (H^+^) from the protonated ε-amino group of Lys74, stabilising the PLP ring in its aromatic state (Figure 5, reaction step 8).

The deprotonated ε-amino group of Lys74 is now ready to reform the internal aldimine with PLP through an analogous process of transamination (steps 1–4) with Lys74 in place of OSS. The ε-amino group of Lys74 forms an imine linkage with PLP, releasing the product L-cysteine (Figure 5, reaction step 9).

### Kinetic Parameters of AfuCys1a and Small Molecule Inhibition

The reaction mechanism of Cys1a enzymes is most consistent with the conserved PLP-dependent ping-pong β-substitution mechanism described for CysK, where divergence in kinetic parameters of Cys1a arises primarily through altered catalytic optimisation rather than major changes in substrate recognition. Previous work studying SpCys1a has indicated that Cys1a enzymes preferentially catalyse the conversion of L-OSS and a reactive sulphur species (RSS) to L-cysteine and succinate (Bastard et al., 2017; X. Zhang et al., 2021). However, the physiologically relevant RSS substrate or substrates are currently unknown, meaning accurate evaluation of this specific activity would not be possible.

Previous work also demonstrated that SpCys1a can perform the canonical CysK reaction, which utilises L-OAS as a substrate with H₂S for the synthesis of L-cysteine (4.5 × 10³ M⁻¹ s⁻¹), but is almost three times more efficient when L-OAS is substituted for L-OSS (1.2 × 10⁴ M⁻¹ s⁻¹) (Bastard et al., 2017). This suggests that L-OAS is the preferred physiological substrate for SpCys1a activity, although it may not be strictly exclusive, as the enzyme can still form the aminoacrylate intermediate and complete the catalytic cycle with L-OAS, albeit with a moderate (∼3-fold) reduction in catalytic efficiency. As Cys1a enzymes can perform the OASS reaction, this could provide an alternative method to assess catalytic activity of AfuCys1a, and would also allow comparison with several related cysteine synthases (Gaitonde, 1967; Pederick et al., 2024).

The activity of AfuCys1a performing the OASS reaction was measured by the cysteine ninhydrin assay, and data was fit using a substrate inhibition model (Figure 6). The model indicated substrate inhibition at concentrations above 6 mM, suggesting there is an optimal substrate range under native conditions which could reflect concentration of available substrate or regulatory mechanisms that prevent excessive product formation (Reed et al., 2010; Y. Zhang et al., 2022). In comparison to SpCys1a in the OASS reaction (Bastard et al., 2017), AfuCys1a exhibited approximately 5-fold lower substrate affinity for L-OAS (*K_m_* = 2.59 ± 0.76 mM), 4-fold higher turnover rate (*k_cat_* = 5.71 ± 1.12 s⁻¹), and 2-fold lower catalytic efficiency (*k_cat_* / *K_m_* = 2.20 × 10³ M⁻¹s⁻¹). Although these values are not vastly different, enzymes with the same predicted catalytic activity are expected to demonstrate highly similar catalytic behaviour. However, several variations in assay conditions between the methods used here and the work studying SpCys1a could reasonably account for this degree of variation, including differences in detection methods, pH, and the use of additional enzymes (McDonald & Tipton, 2022). To better interpret the variation in kinetic values between AfuCys1a and SpCys1a, comparison of was also made with homologues that preferentially catalyse the OASS reaction, and to reduce variation due to assay conditions, values were only included that were measured by the acid ninhydrin assay (Table 2).

**Figure 6.**
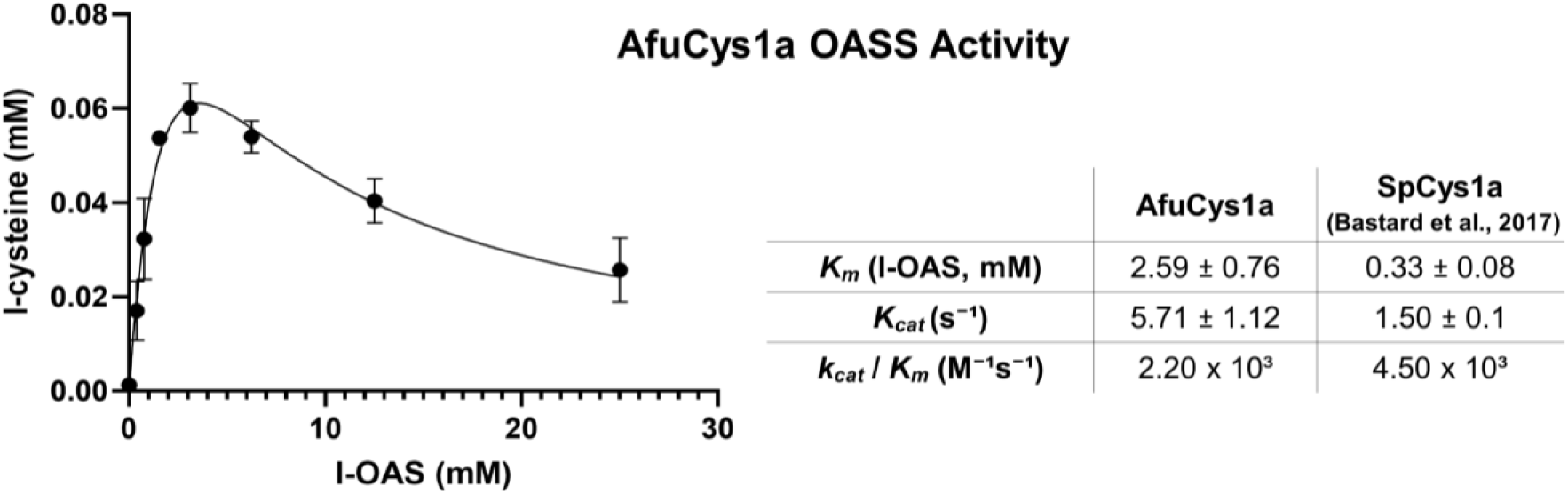
OASS Activity of AfuCys1a in the Cysteine Ninhydrin Assay. The OASS reaction was measured by production of L-cysteine from H2S and varying concentrations of L-OAS and data was fitted to a substrate inhibition model (Eqn. 1).

**Table 2.**
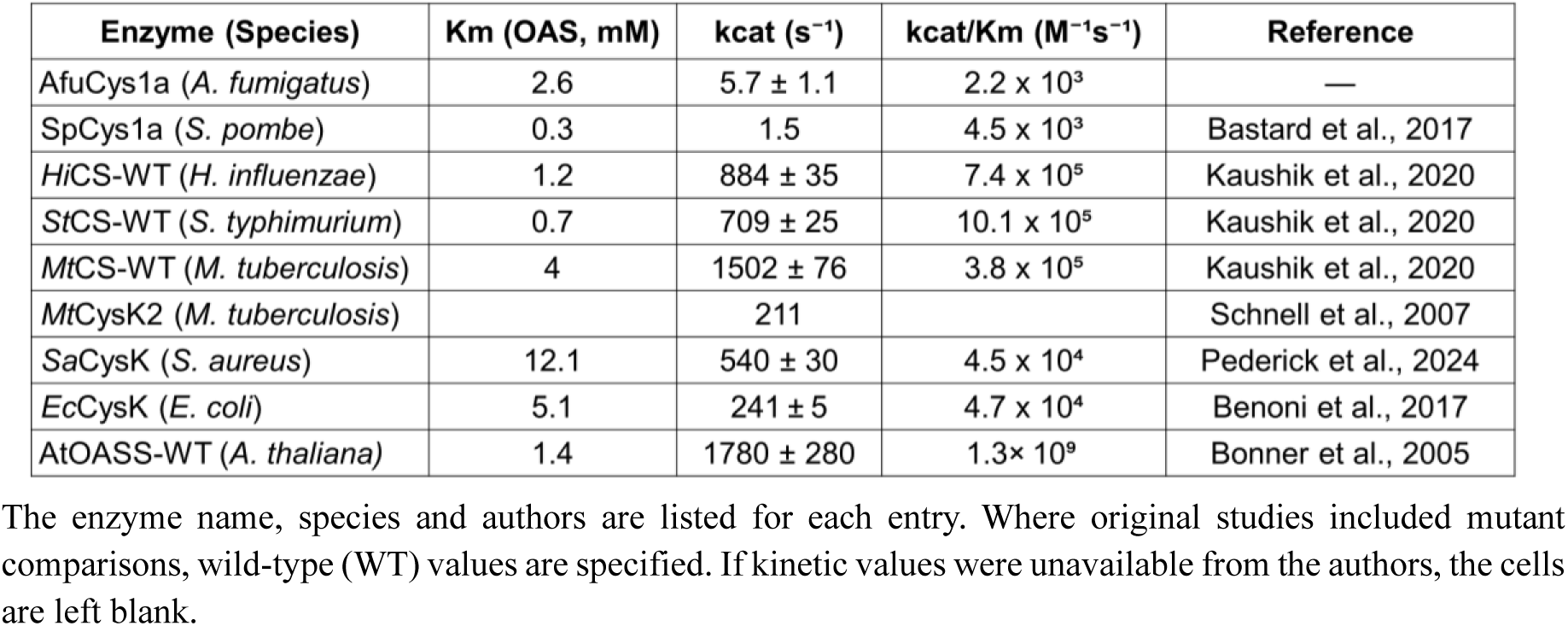
Kinetic values of AfuCys1a and SpCys1a compared with dedicated OASS enzymes The enzyme name, species and authors are listed for each entry. Where original studies included mutant comparisons, wild-type (WT) values are specified. If kinetic values were unavailable from the authors, the cells are left blank.

| Enzyme (Species) | K<sub>m</sub> (OAS, mM) | k<sub>cat</sub> (s<sup>-1</sup>) | k<sub>cat</sub>/K<sub>m</sub> (M<sup>-1</sup>s<sup>-1</sup>) | Reference |
| --- | --- | --- | --- | --- |
| AfuCys1a (*A. fumigatus*) | 2.6 | 5.7 ± 1.1 | 2.2 × 10<sup>3</sup> | — |
| SpCys1a (*S. pombe*) | 0.3 | 1.5 | 4.5 × 10<sup>3</sup> | Bastard et al., 2017 |
| HiCS-WT (*H. influenzae*) | 1.2 | 884 ± 35 | 7.4 × 10<sup>5</sup> | Kaushik et al., 2020 |
| StCS-WT (*S. typhimurium*) | 0.7 | 709 ± 25 | 10.1 × 10<sup>5</sup> | Kaushik et al., 2020 |
| MtCS-WT (*M. tuberculosis*) | 4 | 1502 ± 76 | 3.8 × 10<sup>5</sup> | Kaushik et al., 2020 |
| MtCysK2 (*M. tuberculosis*) |  | 211 |  | Schnell et al., 2007 |
| SaCysK (*S. aureus*) | 12.1 | 540 ± 30 | 4.5 × 10<sup>4</sup> | Pederick et al., 2024 |
| EcCysK (*E. coli*) | 5.1 | 241 ± 5 | 4.7 × 10<sup>4</sup> | Benoni et al., 2017 |
| AtOASS-WT (*A. thaliana*) | 1.4 | 1780 ± 280 | 1.3 × 10<sup>9</sup> | Bonner et al., 2005 |

A substantial range in enzyme activity can be observed between the dedicated OASS enzymes (Table 2), likely reflecting true functional divergence within this group (Kaushik et al., 2021; Kumaran et al., 2009). Despite this range, clear differences in enzyme activity are apparent between Cys1a and OASS enzymes. The bacterial OASS enzymes were approximately 20- to 460-fold more efficient (M⁻¹s⁻¹) at catalysing the OASS reaction than AfuCys1a, and the eukaryotic OASS from *Arabidopsis thaliana* was strikingly more efficient with *k_cat_* / *K_m_* = 1.3× 10⁹ M⁻¹s⁻¹ (Bonner et al., 2005). Similarly, the turnover rate for L-OAS was vastly different between Cys1a and OASS enzymes, with Cys1a enzymes demonstrating 40- to 1000-fold faster turnover of L-OAS (*k_cat_* = 1.5–5.7 s⁻¹) in comparison to dedicated OASS enzymes (241–1780 s⁻¹). This implies that the first half-reaction, encompassing external aldimine formation and β-elimination, remains somewhat permissive and is only moderately tuned towards L-OSS in Cys1a enzymes (Catazaro et al., 2014; Davidi et al., 2016; Rabeh & Cook, 2004).

Further mechanistic insight is also provided by prior studies of SpCys1a that determined kinetic parameters for the second half-reaction, involving nucleophilic attack by a sulphur donor (second substrate) of the α-aminoacrylate intermediate formed from the first substrate (X. Zhang et al., 2021). This work showed that, despite their distinct chemical properties, both GSSH and H₂S are utilised by SpCys1a with comparable catalytic efficiencies (*k_cat_* / *K_m_* = 5.82 × 10³ and 4.03 × 10³ M⁻¹s⁻¹, respectively), whereas thiosulfate is processed substantially less efficiently (*k_cat_* / *K_m_* = 2.81 × 10² M⁻¹ s⁻¹). The similar efficiencies observed for GSSH and H₂S indicate relatively limited discrimination between biologically relevant sulphur nucleophiles, suggesting that SpCys1a could accommodate various sulphur donors during the second half-reaction. This behaviour contrasts with the greater substrate specificity typically associated with OASS enzymes, which are optimised for the canonical L-OAS + H₂S reaction, and instead resembles the broader sulphur donor tolerance reported for CBS-like enzymes involved in sulphur metabolism (Conter et al., 2020; Singh et al., 2009). Such substrate flexibility may reflect adaptation to dynamic physiological conditions, such as fluctuating sulphur availability within mitochondria. Given that mitochondrial sulfide:quinone oxidoreductase (SQR) is proposed to generate the RSS utilised by SpCys1a, promiscuity of the second substrate may enable sustained cysteine biosynthesis across variable mitochondrial sulphur environments (X. Zhang et al., 2021). In this context, there could be several diverse mitochondrial RSS that could function as a more efficient nucleophile within the active site of Cys1a enzymes (Domán et al., 2023; Kolluru et al., 2020; Sawa et al., 2020). This raises the possibility that the physiologically relevant substrate combination may support higher catalytic efficiencies than those tested to date, and further experimental validation would be an important next step in understanding the functional role of AfuCys1a (Robinson, 2015).

Following kinetic characterisation of AfuCys1a, small molecules were screened for their potential inhibitory effects and measured by the cysteine ninhydrin assay. This screening served two intended purposes: first, to help identify LigA and LigB, and second, to provide a foundation for future inhibitor design. LigA and LigB most likely derived from *E. coli* during protein expression and were not deliberately introduced in high concentrations at any stage. This indicates the cyclic molecule would have high affinity for AfuCys1a and should cause enzyme inhibition by competitive binding. Furthermore, LigA and LigB cause disruption to the second substrate loop in a comparable manner to a designed inhibitor (compound 5) in the structural homologue *Mt*CysM (Brunner et al., 2016). Therefore, compounds that do not cause inhibitory effects against AfuCys1a would be excluded as the identity of LigA and LigB, while the structure of LigA and LigB could also provide a basis for small molecule inhibitor selection in the context of future drug development (Koes & Camacho, 2012).

Initially, several molecules known to exhibit inhibitory effects against related enzymes were screened against AfuCys1a, including L-cycloserine, D-cycloserine, L-methionine and L-cystathionine, but showed no inhibition (Aitken & Kirsch, 2003; Franco et al., 2017; Kobylarz et al., 2016; Zhuang et al., 2025). Next, compounds were screened that both resemble the structure of the cyclic molecule and can also be found naturally within *E. coli* cells. These molecules included L-proline, L-pipecolic acid and pyroglutamic acid, but also caused no inhibition (Gold et al., 2022; Guo et al., 2013). Following this, compounds including 2-picolinic acid, 3-picolinic acid (nicotinic acid), 3-methylpyridine-2-carboxylic acid, and 6-hydroxypicolinic acid were screened for inhibition. Compounds were then tested primarily as foundational inhibitors and included 2-picolinic acid, 3-picolinic acid (nicotinic acid), 3-methylpyridine-2-carboxylic acid, and 6-hydroxypicolinic acid. These compounds were not considered for the identity of the LigA and LigB as picolinic acid-based compounds lack the required flexibility within the pyridine ring to reasonably fit the electron density for the cyclic molecule and these are also not considered endogenous metabolites of *E. coli* (Guo et al., 2013). All picolinic acid-based compounds demonstrated similar weak inhibitory activity with preliminary IC₅₀ values ranging from 5 to 15 mM. The assay was repeated three times for 2-picolinic acid providing an IC₅₀ = 11.62 ± 1.85 mM, demonstrating that picolinic acid could be used as a base for the development more potent inhibitors against AfuCys1a (Figure 7).

**Figure 7.**
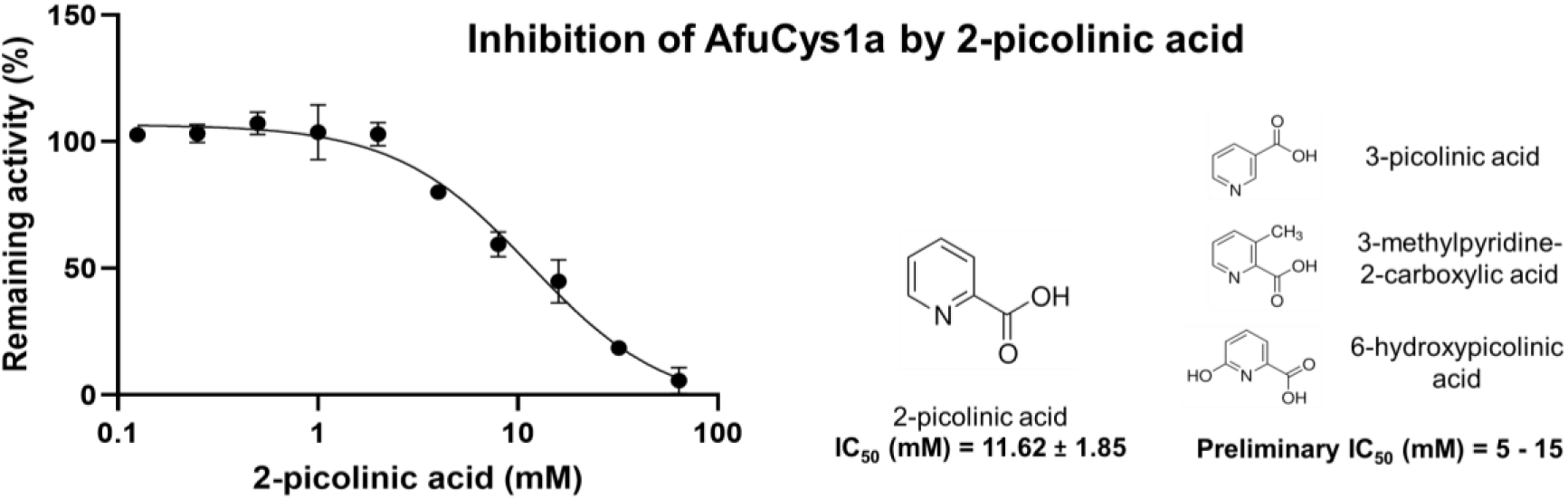
**Inhibition of AfuCys1a by 2-picolinic acid Measured by the Cysteine Ninhydrin Assay**

Electron density for LigA and LigB initially suggested that L-proline would be a reasonable candidate for the identity of the cyclic molecule, but this amino acid has now been excluded by inhibitory assays. Instead, another promising candidate would be the non-proteinogenic amino acid thiazolidine-4-carboxylate (thioproline) which comprises a carboxyl group, and a five-membered ring that contains both a nitrogen and a sulphur atom (S. Kim et al., 2025). Thioproline can also be formed naturally in *E. coli* cells through a non-enzymatic reaction between cysteine and formaldehyde (Kamps et al., 2019; Patterson et al., 2020). The incidental but natural formation of thioproline in is further supported by the identification of enzymes such as proline dehydrogenase and pyrroline-5-carboxylate reductase that may participate in the oxidative metabolism of thioproline within *E. coli* cells (Mao et al., 2021). Moreover, thiazolidine products can form within enzyme active sites or in an enzyme independent manner through a reaction between PLP and L- or D-cysteine and altered by the presence of additional compounds or elements (Kobylarz et al., 2016; Mulay et al., 2023; Nakamura & Fujishiro, 2025). This suggests a mechanism whereby thioproline could be trapped in the enzyme active site through related interactions with PLP as a substrate or product analogue inhibitor of AfuCys1a.

Unfortunately, thioproline could not be confirmed nor excluded as the identity of the cyclic molecule using the cysteine ninhydrin assay. Under assay conditions, thioproline breaks down into cysteine, significantly affecting the amount of product measured using this method. The identity of LigA and LigB or the presence of thioproline in purified AfuCys1a or crystallised protein also could not be determined by mass spectrometry. Results were inconclusive, potentially reflecting limitations in sample preparation, such as insufficient separation from AfuCys1a, instability or loss of the cyclic compound during handling, or degradation into unidentified products. Instead, alternative avenues may need to be explored if LigA and LigB are to be experimentally validated. Despite the unresolved identity of the cyclic molecule, these results demonstrate that structural insights can still provide a valuable foundation for guiding small molecule inhibitor selection as seen through inhibition of AfuCys1a by picolinic acid.

### Identification of Key Residues Involved in Catalytic Specificity of AfuCys1 and Structural Homologues

Careful structural and sequence alignment of AfuCys1a with structural homologues highlighted several residues and structures that could be linked to catalytic specificity. Many close enzyme homologues will perform related but distinct catalytic activity, and consequently share conserved active site residues that are associated with non-specific functional roles across this family (Bartlett et al., 2003; Jack et al., 2016; Ribeiro et al., 2020; Riziotis et al., 2023). These residues are often essential for the core catalytic mechanism, maintaining general active site architecture, or binding cofactors, and are thus subject to strong evolutionary constraints (Jack et al., 2016; MacDonald & Berger, 2014; Ribeiro et al., 2020; Yu et al., 2005). In contrast, active site residues that play a role in determining substrate or catalytic specificity are often uniquely conserved among orthologues with the same enzyme function (Jack et al., 2016; McMurrough et al., 2014; Ribeiro et al., 2020).

The enzyme activity of each homologue, including their preferred substrates and reaction products, are summarised in Table 3. Enzymes used for structural and/or sequence alignments include *Saccharothrix mutabilis* CmnB, *Staphylococcus aureus* SbnA, AfuCys1a, SpCys1a, *Homo sapien* CBS (HsCBS), *Sacharomyces cerevisieae* CBS (ScCBS), *Staphylococcus aureus* OAS-dependent CBS (SaOCBS), *Lactobacillus plantarum* OCBS (LpOCBS), *Bacillus anthracis* OCBS (BaOCBS), *Pseudomonas aeruginosa* CysM (PaCysM), *Salmonella Typhimurium* CysM (StCysM), *Escherichia coli* CysM (EcCysM), *Fusobacterium nucleatum* F1220 lanthionine synthase (FnLS), *Staphylococcus aureus* CysK (SaCysK), *Salmonella Typhimurium* CysK (StCysK), and *Mycobacterium tuberculosis* (MtCysK).

**Table 3.**
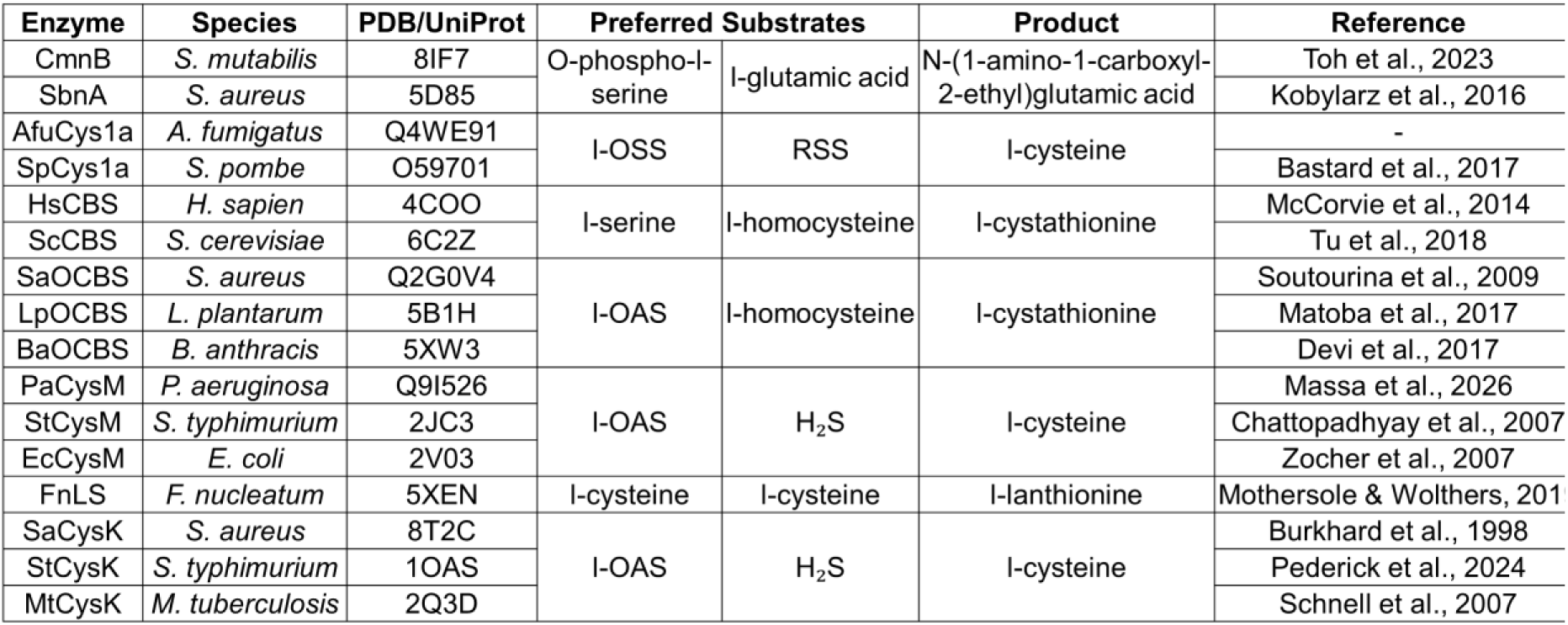
Structural Homologues of AfuCys1a with Distinct Catalytic Activity.

Sequence alignments were complemented by structural alignment to confirm the spatial positioning of each aligned residue. At least two enzymes are compared for each reaction included, except for FnLS which is the sole characterised lanthionine synthase. However, the sequence and structure of FnLS is included here as mutational analysis have identified distinct active site structures and residues that play a role in catalytic specificity (Mothersole et al., 2020; Mothersole & Wolthers, 2019). Similarly, there are no published structures for SpCys1a, SaOCBS, and PaCysM. For each, AlphaFold models were produced and aligned with known structures. This predicted near identical overall fold and conserved active site architecture with enzymes that perform the same reaction.

Analysis indicated that residues Ser129, Lys132, Gln176, Arg340 and Tyr341 are moderately to highly conserved within this family, suggesting a broader functional role rather than key determinants of catalytic specificity (Jack et al., 2016; McMurrough et al., 2014; Ribeiro et al., 2020). Similarly, the majority of the PLP binding sites, the Asparagine Loop, and a small part of the second substrate loop are moderately conserved (Liang et al., 2019; Schnell et al., 2015). However, small sequence variations at these sites have been associated with distinct enzymatic activity. For instance, the Asparagine Loop adopts the sequence motif TAGNT in Cys1a and OCBS enzymes, while this sequence is TSGNT for CysK, CysM, and CBS. Similarly, the PLP phosphate binding site is GTGGT for Cys1a, CysK, and CBS enzymes, whereas this motif is generally GSGGT for OCBS, and GTTGT in CysM. Although most residues within these motifs are highly conserved, substitution at position Ala102 in the Asparagine Loop and variation in the PLP phosphate binding site have been associated with specific catalytic mechanisms. Previous research of serine racemase showed that mutation of serine within the Asparagine Loop (corresponding to Ala102 in AfuCys1a) negatively effects racemisation activity, but β-elimination activity is unaffected (Koulouris et al., 2022; Nelson et al., 2017; Wang et al., 2012). Another study of SbnA, indicated that mutation of first serine within the PLP phosphate binding site motif STTGS (which corresponds to Gly210 in AfuCys1a) may be important for substrate recognition and turnover (Kobylarz et al., 2016). It is likely that variations in both the PLP-phosphate binding motif and position Ala102 (AfuCys1a) within the Asparagine Loop play a role in determination of more general catalytic mechanisms within this protein family, such as preference for racemisation or β-elimination, but variations are unlikely to be highly involved in catalytic specificity.

Instead, key determinants of specificity in AfuCys1a may comprise two active site residues (Thr127 and Gln128), three residues within the second substrate loop (Ser259, Thr261 and Glu262), and an area that forms a clef-like channel or ligand binding pocket (Figure 8). This pocket was initially observed to bind a molecule of glycerol (PDB: GOL) in the holo state of AfuCys1a and is primarily formed by a loop between beta-strand 5 and helix 6, comprising active site residues Phe151–Asn158 and residues Gln256–Arg258 (within the second substrate loop). This pocket is also formed through residues Gln176 and Phe177, though these are highly to moderately conserved within this enzyme family and therefore not suggested here to be key determinants of specificity. A ligand binding pocket an analogous location to AfuCys1a has been described previously in SbnA and FnLS, and is suggested to play a role in specificity due to the presence of a substrate analogue and biologically relevant enzyme product, respectively (Kobylarz et al., 2016; Mothersole & Wolthers, 2019). Additionally, the structures of HsCBS (PDB:4COO) and Human serine racemase (HsSR) (PDB: 6ZSP) contain a molecule of ethylene glycol (PDB: EDO) in an analogous location, potentially suggesting conservation of this ligand binding pocket between homologues, even in enzymes with low sequence identity such as HsSR (Koulouris et al., 2022). Comparing the ligand binding interactions between the structures of SbnA, LnFS, AfuCys1a (holo), and HsCBS at this site could inform a hypothetical mechanism of substrate binding.

**Figure 8.**
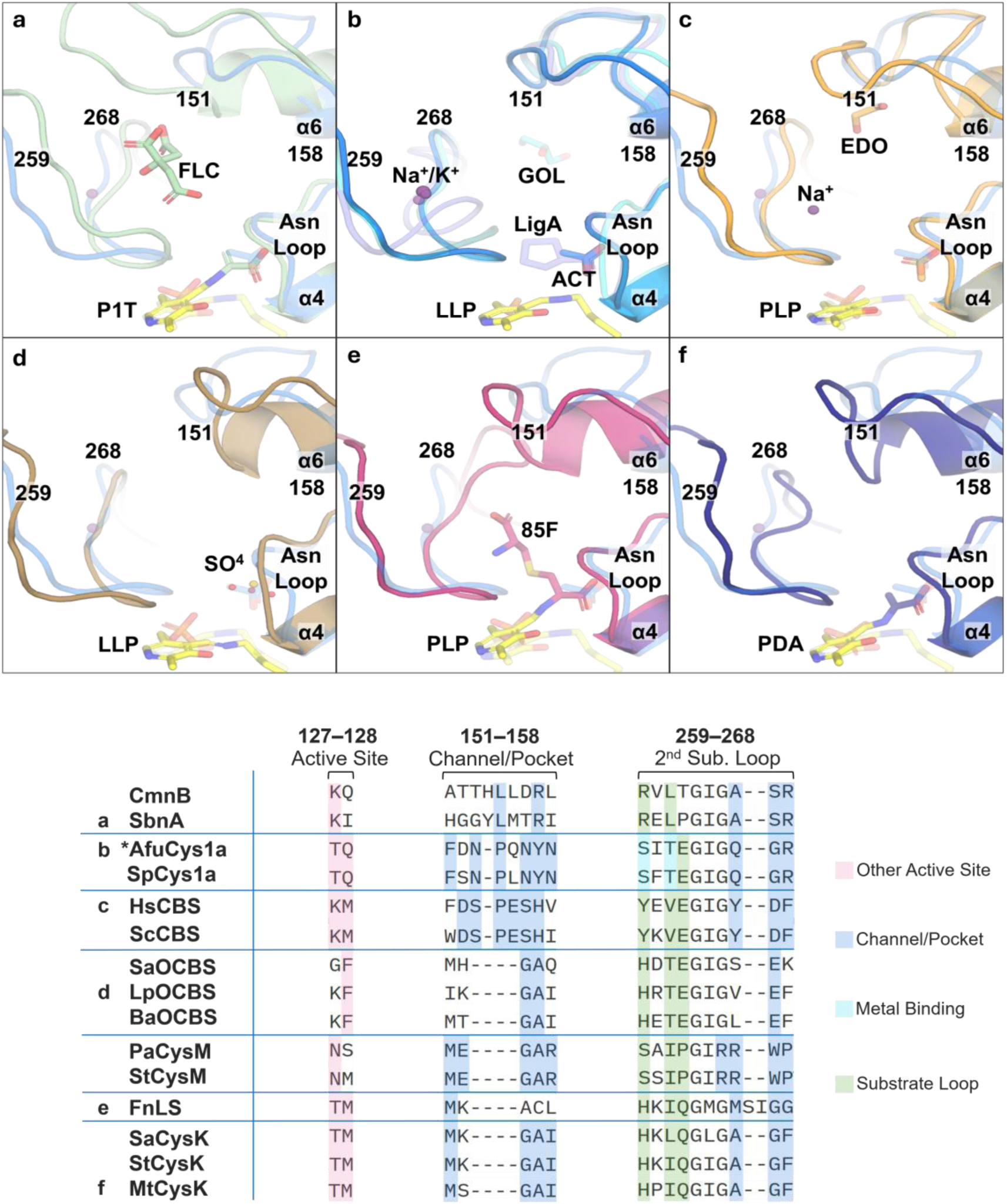
Key Residues and Features Associated with Catalytic Specificity in AfuCys1a and Structural Homologues. Panels **a–f** correspond to enzymes marked **a–f** in the multiple sequence alignment. Each structure is superimposed with the acetate-bound state of AfuCys1a. For AfuCys1a (panel **b**), the acetate-bound state (dark blue) is superimposed with the three other states of AfuCys1a (transparent), including holo (cyan), LigA-bound (slate), and LigB-bound (teal). The PDB codes for each structure are included in Table 3.

It should be noted that the enzymes HsCBS and ScCBS possess additional CBS-specific regulatory domains that are absent from the other enzymes examined here. These domains are linked to modulation of catalytic activity and may influence the conformational state and accessibility of the catalytic core (McCorvie et al., 2014; Tu et al., 2018). Consequently, differences in catalytic behaviour between CBS enzymes and proteins lacking these regulatory elements may not solely reflect the effect of specific residues or variations within the active site architecture but instead may also arise from regulatory mechanisms associated with these auxiliary domains.

A link between residues that form the ligand binding pocket and substrate specificity can also be supported by previous mutational studies. For CBS and OCBS, prior work demonstrated that the presence of a tyrosine at the position corresponding to Gln266 in AfuCys1a (Figure 8) was strongly associated with substrate specificity for L-serine in CBS enzymes (Conter et al., 2023; Devi et al., 2019; Lodha et al., 2009). For SbnA, the residues Arg132 and Tyr152 correspond to Tyr157 and Phe177 in AfuCys1a (Figure 8). Mutation of SbnA R132A and Y152F indicated residues at these positions may be important for substrate recognition and turnover (Kobylarz et al., 2016). Additionally, Arg132 of SbnA (Tyr157 for AfuCys1a) interacted with the citrate molecule, which resembles the enzymes preferred substrate, L-glutamate, associating this residue with substrate binding. Similarly, SbnA residues Lys100, and Arg224 also interact with a citrate molecule and correspond to AfuCys1a positions Thr127 and Ser259 (Figure 8). For CysK enzymes, mutation of methionine at position Met120 (*Haemophilus influenzae* CysK numbering) leads to a marked decrease in substrate binding and catalytic efficiency. Methionine at this position is described non-catalytic residue located approximately 20 Å from the reaction centre and corresponds to Phe151 in AfuCys1a (Figure 8), also located within the ligand binding pocket (Kaushik et al., 2021).

Studies of FnLS also linked the residues MSIG (residues 223–226) to catalytic specificity (Mothersole & Wolthers, 2019). This sequence corresponds to Gln266–Gly267 in AfuCys1a (Figure 8) but also contains two additional residues, Ser224 and Ile225, which are absent in other homologues. These additional residues in FnLS result in an extended part of second substrate loop which corresponds to the ligand binding pocket. Residues in this extended loop are linked to specificity for L-lanthionine, as they interact with the lanthionine-PLP external aldimine and undergo conformational changes upon ligand binding. Mutational analysis further shows that substitution of Ser224 with alanine (S224A) increases turnover sevenfold but reduces specificity for lanthionine over cystathionine, illustrating a trade-off between catalytic rate and specificity at this site (Mothersole et al., 2020). The FnLS residues Met223 and Gly226 correspond to AfuCys1a Gln266 and Gly267 within the second substrate loop and comprise part of ligand binding pocket.

The extended loop of FnLS exemplifies how shape and structure as well as specific amino acids can affect enzyme catalysis. However, several mutational studies have struggled to pinpoint individual residues, or even combinations of residues, as strong determinants of enzyme specificity. This is because catalytic specificity is not often entirely dictated by the chemical properties of one residue alone. Instead, specificity is the result of structural and spatial geometry in combination with local chemical environments. In addition to the extended loop FnLS, alignment of the ligand pocket highlights distinct structures between enzymes that perform different functions, including the length of the loop that corresponds to Phe151–Asn158 in AfuCys1a and the extension of helix 6.

The active site of AfuCys1a appears spatially organised to stabilise larger RSS substrates, which may hinder optimal positioning of the smaller H₂S molecule for efficient nucleophilic attack (Catazaro et al., 2014). To test this, docking studies with the predicted substrates L-OSS-PLP and GSS^-^ yielded multiple poses for each ligand in similar locations within the active site. These poses formed hydrogen bonds with, or were spatially oriented to interact with, residues hypothesised to govern substrate specificity and catalytic activity in AfuCys1a. The final poses presented here exhibited Edock scores between −40.5 and −49.1 kcal mol⁻¹ and were positioned in regions consistent with the proposed reaction mechanism for these substrates (Figure 9). Notably, one previous pose of GSS^-^ within the active site of AfuCys1a also showed hydrogen bond formation with active site residues Thr127 and Gln128 (Edock = −40.48 kcal mol⁻¹), supporting their previous implication in specificity. Collectively, these docking results are consistent with L-OSS and GSS^-^ being relevant substrates and support the hypothesis that the identified residues (Figure 8) contribute to catalytic specificity in AfuCys1a.

**Figure 9.**
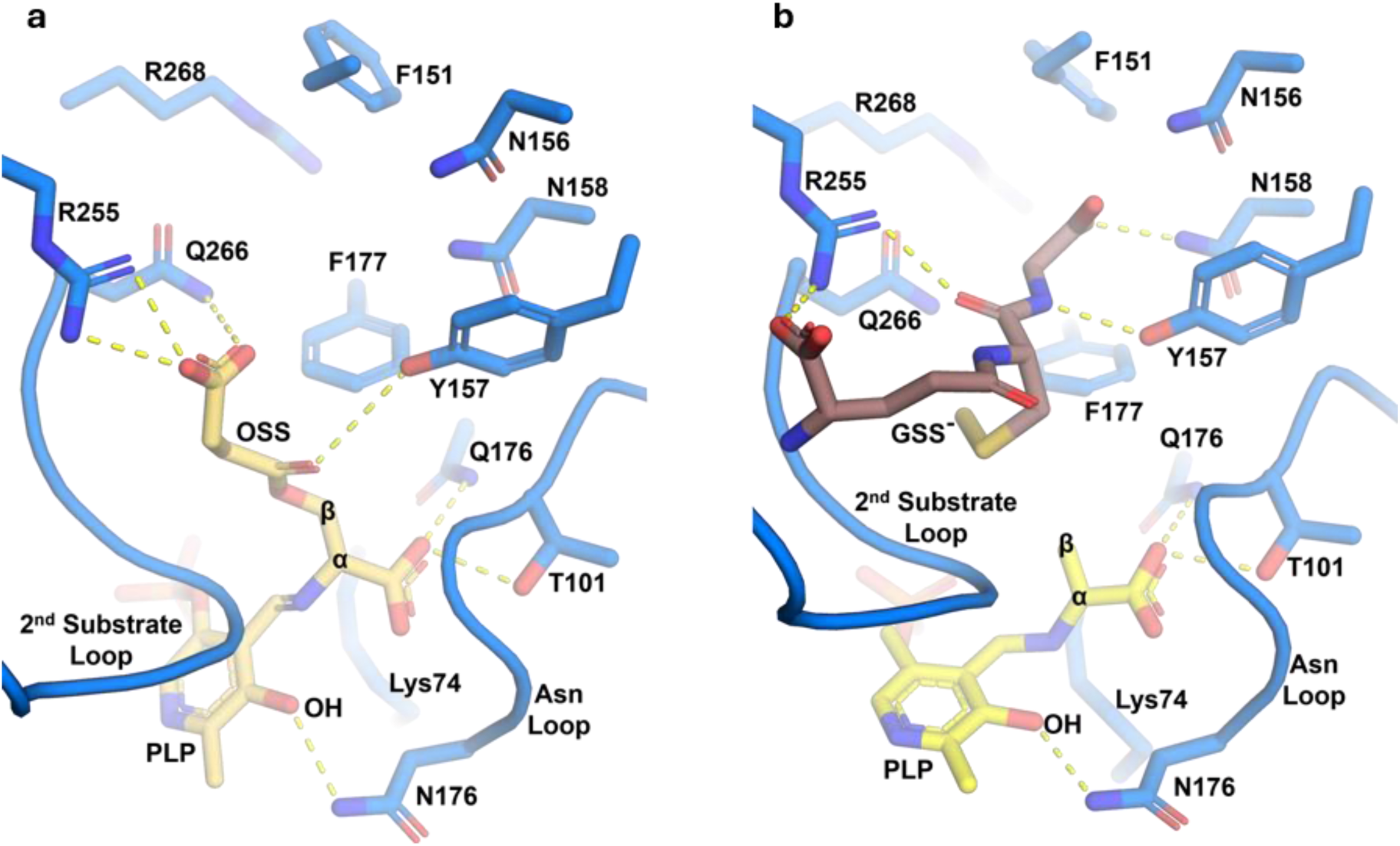
Substrate Docking of L-OSS and GSS^-^ in the acetate-bound state of AfuCys1a. **a** Pose of PLP-L-OSS external aldimine complex in the active site of AfuCys1a. **b** Pose GSS^-^ within the active site of AfuCys1a, positioned in a manner that would allow nucleophilic attack of the β-carbon of the PLP α-aminoacrylate.

## Conclusion

This study provides the first detailed structural and biochemical characterisation of AfuCys1a, a member of the recently described Cys1a family of PLP-dependent enzymes. High-resolution crystal structures revealed a conserved PLP-dependent fold type II architecture and identified key active site features, including the PLP-binding site, Asparagine Loop, second substrate loop, and a putative ligand binding pocket. Comparison of four ligand-bound states demonstrated that ligand binding to the Asparagine Loop promotes a closed active site conformation associated with alignment of catalytic residues, while disruption caused by the unidentified ligands LigA and LigB is partially mitigated by metal coordination and hydrogen-bonding interactions of Glu262. These findings highlight the importance of active site dynamics in regulating enzyme function and provide a structural framework for understanding catalysis within the Cys1a family.

Kinetic analysis demonstrated that AfuCys1a catalyses the OASS reaction using L-OAS and H₂S, but with substantially lower catalytic efficiency than dedicated OASS enzymes. Comparison with previously characterised SpCys1a supports the hypothesis that Cys1a enzymes retain the ability to utilise L-OAS while being adapted for alternative physiological substrates, most likely L-OSS and a reactive sulphur species. Structural comparisons, sequence conservation analyses, and molecular docking further identified residues and active site regions that may contribute to catalytic specificity, such as residues within the second substrate loop, the adjacent ligand binding pocket, as well as the active site residues Thr127 and Gln128. Together, these observations support a model in which substrate specificity is determined by a combination of active site chemistry, loop dynamics, and three-dimensional architecture rather than by a small number of catalytic residues alone.

Although the identity of LigA and LigB remains unresolved, their repeated observation in the active site, apparent strong binding affinity, and disruption of catalytically important regions suggest that they may represent inhibitory molecules. Consistent with this interpretation, small-molecule screening identified picolinic acid derivatives as weak inhibitors of AfuCys1a and demonstrated that structural information can be used to guide inhibitor selection. Given the importance of sulphur metabolism for fungal survival and virulence, these findings establish a foundation for the rational design of compounds targeting Cys1a enzymes.

Overall, this work advances current understanding of this enzyme family by integrating structural biology, enzymology, inhibition studies, and comparative analyses. Analysis of kinetic parameters support the emerging model that Cys1a enzymes participate in sulphur metabolism and may contribute to the intersection of sulphur homeostasis, mitochondrial function, and cellular signalling by permissive use of RSS. Future studies aimed at identifying the physiological substrates of AfuCys1a, experimentally validating the proposed reaction mechanism, and resolving the identity of LigA and LigB will be essential for fully defining the biological role of this unusual class of cysteine synthases.

## Acknowledgements

I would like to acknowledge that this research was undertaken in part using the MX1 and MX2 beamlines at the Australian Synchrotron, part of ANSTO, and made use of the Australian Cancer Research Foundation (ACRF) detector. Acquiring the protein structures presented here would not have been possible without this research collaboration. Dr Blagojce Jovcevski is supported by a Hospital Research Foundation Research Fellowship (2023/QA25313).

